## Supplemental Material 3 for "Messinian vegetation and climate of the intermontane Florina-Ptolemais-Servia Basin, NW Greece inferred from palaeobotanical data: How well do plant fossils reflect past environments?" (from another version of this preprint: version 1, 2019-11-25)

**Supplementary Material S3:** Systematic palaeobotany and descriptions of palynomorphs

from the plant fossil bearing strata of Vegora (sample S115992)

The systematic palaeobotany section starts with algae and is followed by fern and fern allies, gymnosperms, and angiosperms. Angiosperm classification and author names of orders and families follow APG IV [1]. LM and SEM in the descriptions refers to light microscopy and scanning electron microscopy observations, respectively.

(a) Algae

Family Botryococcaceae
Genus *Botryococcus* Kürtzing
*Botryococcus* sp. (Fig. 3a)

Remarks: Similar structures are produced by extant *Botryococcus brauni* Kürtzing *Botryococcus* is commonly found in freshwater but can thrive also in brackish conditions.

Family Zygnemataceae Kützning
Genus *Spirogyra* Link in C.G. Nees/ *Ovoidites* R.Potonié emend. Krutzsch
*Spirogyra* sp. 1/ *Ovoidites elongatus* (Hunger) Krutzsch (Fig. 3b)

Description: Aplanospore or zygospore, outline elliptic to broadly ovoidal, palynomorph size large, length of axis perpendicular to fissure 70–100 µm (LM); mesospore 1–1.5 μm thick (LM); sculpturing psilate (LM, SEM), if ruptured gap spanning from pole to pole.

Remarks: *Ovoidites elongatus* is commonly associated with *Spirogyra*, indicating shallow, stagnant, oxygen-rich open freshwaters and lake margins [2].

Genus *Cycloovoidites* Krutzsch & Pacltová
*Spirogyra* sp. 2/*Cycloovoidites cyclus* (Krutzsch) Krutzsch & Pacltová (Fig. 3c)

Description: Aplanospore or zygospore, outline circular, palynomorph size large to very large, diameter 90–110 µm (LM); mesospore 2–2.4 μm thick (LM); sculpturing rugulate, verrucate (LM, SEM).

Remarks: *Cycloovoidites cyclus* is commonly associated with *Spirogyra* [2, 3].

(b) Fern and Fern allies

Order Osmundales Bromhead
Family Osmundaceae Marinov
Genus *Osmunda* L.
*Osmunda* sp. (Figs 3d–e)

Description: Spore, monad, shape spheroidal to oblate, amb circular; spore size medium to large, equatorial diameter 30–65 μm (LM, SEM); exospore 1–1.5 μm thick (LM), 2–3 µm thick including sculpture elements (LM); trilete, laesurae 2/3 to 3/4 of spore radius; sculpturing bacculate to rugulate (LM), sculpturing present on distal and proximal face.

Remarks: Extant *Osmunda* produces morphologically identical spores [4, 5]. Specimens identified from the Vegora sample fall within the range of the fossil form species *Bacculatisporites major* (Raatz) Krutzsch, *B*. *primarius* (Wolff) P.W.Thomson & Pflug and *B*. *nanus* (Wolff) Krutzsch. [6, 7].

Order Polypodiales Link
Family Pteridaceae E.D.M.Kirchn.
Genus *Cryptogramma* R.Brown/ *Cheilanthes* Sw.
*Cryptogramma* vel *Cheilanthes* sp. (Figs 3f)

Description: Spore, monad, shape oblate, amb triangular; spore size medium, equatorial diameter 35–50 μm (LM); exospore 1–1.5 μm thick (LM), <2.5 µm thick including sculpture elements (LM); trilete, laesurae 2/3 to 3/4 of spore radius; sculpturing verrucate (LM), sculpturing less prominent on proximal face.

Remarks: Extant *Cryptogramma* [5] and *Cheilanthes* [8] produce morphologically similar spores.

Genus *Pteris* L.
*Pteris* sp. (Figs 3g–h)

Description: Spore, monad, shape oblate, amb convex triangular; spore size medium, equatorial diameter 25–30 μm (LM); exospore 0.5–1 μm thick (LM), <3.5 µm thick including sculpture elements (LM); trilete, laesurae 2/3 to 3/4 of spore radius, cingulum present; sculpturing bacculate (LM), sculpturing less prominent on proximal face.

Remarks: Several extant *Pteris* species produce morphologically similar spores, e.g. *P. dentata* Forssk., *P. ensiformis* Burm. [5, 9].

Family Davalliaceae M.R.Schomb./ Polypodiaceae J.Presl & C.Presl
Davalliaceae vel Polypodiaceae sp./ *Verrucatosporites alienus* (R.Potonié) P.W.Thomson & Pflug (Fig. 3i)

Description: Spore, monad, shape oblate, amb elliptic, outline elliptic to renal-shaped in equatorial view; spore size medium to large, equatorial diameter 50–60 μm (LM); exospore 1.5–3.5 µm thick including sculpture elements (LM); monolete, laesurae 1/2 to 2/3 of spore radius; sculpturing verrucate (LM), sculpturing less prominent on proximal face.

Remarks: Morphologically similar spores are found in *Davallia* Sm., *Microgramma* C.Presl, *Pleopeltis* Humb. & Bonpl. ex Willd., and *Polypodium* L. [5].

Incerta sedis
Monolete spore fam. indet. sp. /*Laevigatosporites haardti* (R.Potonié & Venitz) P.W.Thomson & Pflug (Fig. 3j)

Description: Spore, monad, shape oblate, amb elliptic, outline elliptic to renal-shaped in equatorial view; spore size medium, equatorial diameter 25–50 μm (LM); exospore 1–1.5 µm thick (LM); monolete, laesurae 1/2 to 2/3 of spore radius; sculpturing psilate (LM).

Remarks: Morphologically similar monolete and psilate spores are found in several extant fern families (e.g. Aspleniaceae, Davalliaceae, Dryopteridaceae, Gleicheniaceae, Lomariopsidaceae, Oleandraceae, Thelypteridaceae, Vittariaceae, Polypodiaceae, see [5, 7]).

(c) Gymnosperms

Family Cupressaceae Rich. ex Bartling
Papillate Cupressaceae pollen/ *Inaperturopollenites hiatus* (R.Potonié) Thomson & Pflug (Figs 3k–l)

Description: Pollen, monad, spheroidal, outline circular; pollen size medium, diameter 25–50 μm (LM); exine 1–1.5 µm thick (LM); leptoma with papilla, radially split with papilla at the end of rupture; sculpturing scabrate (LM), microverrucate, nanoechinate (SEM), orbiculae present.

Remarks: Split papillate Cupressaceae pollen is generally assigned to *Inaperturopollenites hiatus* [10, 11]. Pollen similar to the figured specimen is produced by extant members of subfamilies Sequoioideae and Taxodioideae [12, 13]. Ruptured pollen of these subfamilies is stenopalynous and lacks distinct characteristics for further determination.

Family Pinaceae Spreng. ex F.Rudolphi
Genus *Abies* Mill.
*Abies* sp. (Fig. 3m)

Description: Pollen, monad, bisaccate, shape oblate, outline elliptical in polar view, pollen size large to very large, diameter 70–120 μm (LM); exine 1.5–2 in cappa region 3–5 µm thick (*Abies*-crest in LM); leptoma; sacci nearly spherical.

Remarks: The figured specimen corresponds (e.g. crest, sacci nearly spherical, size) to extant pollen of *Abies* [14].

Genus *Cathaya* Chun & Kuang
*Cathaya* sp. (Figs 3n–o)

Description: Pollen, monad, bisaccate, shape oblate, outline elliptical in polar view, pollen size large, diameter 60–80 μm (LM); exine 1–1.5 µm thick (LM); leptoma; sacci half-spherical; exine sculpturing nanoechinate (SEM).

Remarks: Nanoechinate exine sculpturing (Fig. 3o) is a characteristic feature of extant *Cathaya* pollen [15,16].

Genus *Cedrus* Trew
*Cedrus* sp. (Fig. 3p)

Description: Pollen, monad, bisaccate, shape oblate, outline elliptical in polar view, pollen size large, diameter 60–80 μm (LM); exine 1–1.5 µm thick, in cappa region <3 µm thick (LM); leptoma; sacci half-spherical, sacci attachment on proximal face thickened.

Remarks: The figured pollen resembles extant *Cedrus libani* A.Rich. and *C. deodora* (Roxb. ex D.Don) G.Don [17].

Genus *Pinus* L.
*Pinus* subgenus *Pinus* L.
*Pinus* subgenus *Pinus* sp. (Fig. 3q)

Description: Pollen, monad, bisaccate, shape oblate, outline elliptical in polar view, pollen size large, diameter 60–80 μm (LM); exine 1–1.5 µm thick (LM); leptoma; sacci nearly spherical, sacci attachment narrow.

Remarks: Pollen of *Pinus* subgenus *Pinus* sp. (diploxylon type) is characterized by narrowly attached and spherical sacci [18].

*Pinus* subgenus *Strobus* Lemmon
*Pinus* subgenus *Strobus* sp. (Fig. 3r)

Description: Pollen, monad, bisaccate, shape oblate, outline elliptical in polar view, pollen size large, diameter 60–80 μm (LM); exine 1–1.5 µm thick (LM); leptoma; sacci half-spherical, sacci attachment broad.

Remarks: Pollen of *Pinus* subgenus *Pinus* sp. (haploxylon type) is characterized by broadly attached, half-spherical sacci and dotted thickenings in the leptoma area [18].

Genus *Tsuga* (Endl.) Carrière
*Tsuga* sp. 1 (Figs 3s–t)

Description: Pollen, monad, monosaccate, shape oblate, outline circular in polar view, elliptic in equatorial view, pollen size medium to large, equatorial diameter 45–60 μm (LM); saccus 3–5 µm wide (LM); leptoma; sculpturing verrucate, rugulate, echinate (LM), echini length 1.5–3 µm, echini equally distributed.

Remarks: Morphologically similar echinate pollen with a relatively narrow saccus is produced by extant *Tsuga dumosa* Eichl. [19].

*Tsuga* sp. 2 (Figs 3u–v)

Description: Pollen, monad, monosaccate, shape oblate, outline circular in polar view, elliptic in equatorial view, pollen size large, equatorial diameter 60–80 μm (LM); saccus 5–10 µm wide (LM); leptoma; sculpturing verrucate, rugulate, echinate (LM), echini length 1–2.5 µm, echini density lower in saccus area.

Remarks: *Tsuga* sp. 2 differs from *Tsuga* sp. 1 by larger size and a broader saccus. Morphologically similar echinate pollen with a relatively broad saccus is produced by extant *Tsuga forrestii* Downie and *T. chinensis* (Franch.) E.Pritz. in Diels [19].

(d) Angiosperms

Order Poales Small
Family Typhaceae Juss.
Genus *Typha* L./ *Tetradomonoporites* Chitaley
*Typha* sp./ *Tetradomonoporites typhoides* Krutzsch (Fig. 4a)

Description: Pollen, permanent tetrad, planar tetrad, monad shape spheroidal to subspheroidal, outline circular, tetrad size medium, monad size small, monad diameter 18–25 μm (LM); porate, porus sunken; sculpturing reticulate (LM, SEM).

Remarks: Within Typhaceae, only *Typha* disperses pollen in permanent tetrads [14, 20].

Family Poaceae Barnhart
Poaceae gen indet. (Figs 4b–c)

Description: Pollen, monad, shape spheroidal, outline circular, pollen size small, monad diameter 20–25 μm (LM); porate, porus annulate; sculpturing scabrate (LM), nanoechinate, nanoechini weakly grouped in areolae (SEM).

Remarks: Areolate exine sculpturing is widely present in several Poaceae subfamilies (e.g. *Poa* spp., *Phleum* spp. [21]).

Order Vitales Juss. ex Bercht. J.Presl
Family Vitaceae Juss.
Genus *Parthenocissus* Planch.
*Parthenocissus* sp. (Figs 4f–g)

Description: Pollen, monad, shape prolate, outline elliptic in equatorial view, pollen size medium, diameter 18–25 μm (LM), polar axis 30–35 µm (LM); tricolporate, endoporus lalongate elliptic; sculpturing reticulate (LM, SEM), lumen size decreasing towards colpi (SEM).

Remarks: Pollen similar to the figured specimen is found in extant e.g. *Parthenocissus sinensis* Diels & Gilg ex Diels or *P. heptaphylla* (Planch.) Britton [22].

Order Rosales Bercht. & Presl
Family Ulmaceae Mirbel
Genus *Ulmus* L./ Zelkova Spach.

*Ulmus* vel *Zelkova* sp. (Fig. 4h)

Description: Pollen, monad, shape oblate, outline weakly circular to polygonal in polar view, pollen size medium, diameter 35–55 μm (LM); porate, in some specimen annulus present; sculpturing rugulate (LM).

Remarks: *Ulmus* and *Zelkova* share overlapping morphological characters [23], the encountered specimens display poor preservation. Both *Ulmus* and *Zelkova* are present in the Vegora fossil fleaf record.

Order Fagales Engler
Family Fagaceae Dumort.
Genus *Fagus* (Fig. 4i)

Description: Pollen, monad, shape spheroidal to subprolate, outline circular in polar view, circular to elliptic in equatorial view, pollen size medium, polar axis 35–50 μm long (LM); tricolporate; sculpturing scabrate (LM), rugulate, fossulate, rugulae often protruding and diverging (SEM).

Remarks: Modern pollen of *Fagus* subgenus *Fagus* displays similar size ranges and colpi length ranging from half to two thirds of the polar axis [24, 25].

Genus *Quercus* L.
*Quercus* subgenus *Cerris* sect. *Cerris* Oerst.
*Quercus* sect. *Cerris* sp. (Figs 4j–k)

Description: Pollen, monad, shape prolate, outline lobate in polar view, elliptic in equatorial view, pollen size medium, polar axis 30–40 μm long (LM), diameter; tricolpate; sculpturing scabrate (LM), mircro- to nanorugulate, fossulate, perforate, pollen surface irregularly covered by tufts and agglomerations of rodlet tufts.

Remarks: Generally, the ectexine of *Quercus* is composed of rod-like (micro) rugulae [24]. Sculpturing consisting of rugulae masked by secondary sporopollenin and tufts or agglomerations of tufts is characteristic for pollen of *Quercus* sect. *Cerris* [24].

*Quercus* subgenus *Cerris* sect. *Ilex* Loudon
*Quercus* sect. *Ilex* sp. (Figs 4l–m)

Description: Pollen, monad, shape prolate, outline lobate in polar view, elliptic in equatorial view, pollen size small to medium, diameter 20–30 μm (LM); tricolpate; sculpturing scabrate (LM), micro- to nanorugulate, fossulate (SEM).

Remarks: Sculpturing consisting of rod-like micro to nanorugulae is characteristic of *Quercus* sect. *Ilex* [24].

*Quercus* subgenus *Quercus* sect. *Quercus* Loudon
*Quercus* sect. *Quercus* sp. (Figs 4n–o)

Description: Pollen, monad, shape prolate to subprolate, outline lobate in polar view, elliptic sub cyclic in equatorial view, pollen size medium, diameter 30–45 μm (LM); tricolpate; sculpturing scabrate (LM), micro- to nanoverrucate, fossulate, perforate, sculpture elements cauliflower-like (SEM).

Remarks: Cauliflower-like exine sculpturing is characteristic for pollen belonging to this section [24].

Subfamily Castaneoideae
Castaneoideae gen. indet. (Figs 4p–q)

Description: Pollen, monad, shape prolate, outline circular in polar view, elliptic in equatorial view, pollen size small, diameter 10–15 μm (LM), polar axis 18–25 µm (LM); tricolporate; sculpturing scabrate (LM), rugulate to microrugulate, perforate (SEM), rugulae covered with secondary striation (microrugulae), perforations encircled by triangular units formed by rugulae (SEM).

Remarks: Within the paraphyletic Castaneoideae pollen morphology is highly stenopalynous with several overlapping morphological characteristics, therefore generic determination is not possible in LM and SEM investigation. Similar exine sculpture (secondary striation on rugulae, triangular units) is present in pollen of extant *Castanopsis sieboldii* (Makino) Hatus. (=*C. cuspidata* var. *sieboldii*) [26].

Family Juglandaceae DC. ex Preleb
Subfamily Juglandoideae
Genus *Carya* Nutt.
*Carya* sp. (Fig. 4r)

Description: Pollen, monad, shape oblate, outline circular to convex triangular in polar view, pollen size medium, diameter 35–50 μm (LM); triporate, pori sunken; sculpturing scabrate (LM), nanoechinate (SEM).

Remarks: Oblate pollen with three pores off-set towards the distal pole is characteristic for this genus [27].

Genus *Platycarya*
*Platycarya* sp. (Fig. 4s)

Description: Pollen, monad, shape oblate, outline circular to convex triangular in polar view, pollen size small, diameter 18–25 μm (LM); triporate, stephanoporate, pori sunken; sculpturing scabrate, (LM), nanoechinate (SEM), pseudocolpi present on distal and proximal face (LM, SEM).

Remarks: *Platycarya* pollen features pseudocolpi on both faces [28].

Subfamily Engelhardioideae Iljinsk.
Engelhardioideae gen. indet. (Fig 4t)

Description: Pollen, monad, shape oblate, outline triangular in polar view, pollen size small, diameter 16–22 μm (LM); triporate, stephanoporate, pori sunken; sculpturing scabrate (LM), nanoechinate (SEM).

Remarks: Within Engelhardioideae pollen morphology is highly stenopalynous (triporate, nanoechinate sculpturing) [27]. The Engelhardioideae fossil record documents a diversity much higher than its extant three to four genera (*Alfaroa* Standl., *Alfaropsis* (Wall.) I.A.Iljinskaja, *Engelhardia* Lesch ex Blume, *Oremunnea* Oerst.) and the presence of extinct linages during the Cenozoic [28].

Family Betulaceae Gray
Genus *Alnus* Mill.
*Alnus* sp. (Fig 4u)

Description: Pollen, monad, shape oblate, outline polygonal in polar view, pollen size medium, diameter 25–40 μm (LM); tetra- to pentaporate, stephanoporate, pori annulate, adjacent pori connected by arci; sculpturing scabrate (LM), nanoechinate, two to three nanoechini grouped on microrugulae/ridges (SEM).

Remarks: Arci spanning between adjacent pores are characteristic of *Alnus* pollen [29].

Genus *Betula* L.
*Betula* sp. (Fig 4v)

Description: Pollen, monad, shape oblate, outline convex triagonal convex to circular in polar view, pollen size medium, diameter 25–40 μm (LM); triporate, pori annulate, annulus

formed by sexine, vestibulum present; sculpturing scabrate (LM), nanoechinate, two to three nanoechini grouped on microrugulae/ridges (SEM).

Remarks: A distinct atrium is a characteristic feature of *Betula* [29].

Genus *Carpinus* L.
*Carpinus* sp. (Fig 4w)

Description: Pollen, monad, shape oblate, outline circular in polar view, pollen size medium, diameter 35–50 μm (LM); tetra- to pentaporate, pori, weak aspis present; sculpturing scabrate (LM), nanoechinate, two to three nanoechini grouped on microrugulae (ridges) (SEM).

Remarks: Oblate pollen with weakly protruding apertures is characteristic of *Carpinus* [29].

Genus *Corylus* L.
*Corylus* sp. (Fig 4x)

Description: Pollen, monad, shape oblate, outline convex triangular in polar view, pollen size medium, diameter 25–40 μm (LM); triporate, pori annulate; sculpturing scabrate (LM), nanoechinate, two to three nanoechini grouped on microrugulae (ridges) (SEM).

Remarks: The absence of a distinct atrium and the convex triangular outline in polar view are characteristic of *Corylus* [29].

Order Malpighiales Juss. ex Bercht. & J.Presl
Family Salicaceae Mirb.
Genus *Salix* L.
*Salix* sp. (Fig. 4y)

Description: Pollen, monad, shape prolate to subprolate, outline elliptic in equatorial view, pollen size small, diameter 20–25 μm (LM); colpate; sculpturing reticulate (LM, SEM), muri wedge shaped (SEM).

Remarks: The figured specimen features wedge-shaped muri in SEM investigation; this is the most common muri type in *Salix* pollen [30].

Order Geraniales Juss. ex Bercht. & J.Presl
Family Geraniaceae Juss.
Genus *Geranium* L.
*Geranium* sp. (Figs 4z–aa)

Description: Pollen, monad, shape spheroidal, outline circular to weakly lobate in polar view, circular in equatorial view, pollen size large, diameter 65–80 μm (LM); tricolpate; sculpturing reticulate, clavate (LM, SEM), “heads” of clavae rugulate (SEM).

Remarks: The figured specimen corresponds (reticulum cristatum, clavae with rugulae and circular heads) to the *Geranium robertianum* group of the *Geranium molle* type of Stafford and Blackmore [31].

Order Sapindales Juss. ex Bercht. & J.Presl
Family Anacardiaceae R.Br.
Genus *Cotinus* Mill.

*Cotinus* sp. (Figs 5a–b)

Description: Pollen, monad, shape prolate, elliptic in equatorial view, pollen size medium, diameter 20–30 μm (LM), polar axis 25–35 µm long (LM); tricolporate, lalongate rhombic endoporus; sculpturing striate (LM), striatoreticulate (SEM), striae in colpus region more densely packed (SEM).

Remarks: The figured specimen displays typical characters (striatoreticulum, rhombic endoporus) of *Cotinus* [32].

Genus *Pistacia* L.
*Pistacia* sp. (Figs 5c–d)

Description: Pollen, monad, shape spheroidal, outline circular in polar view, pollen size medium, diameter 25–35 μm (LM); penta- to hepta pantoporate, pori sunken; sculpturing scabrate (LM), reticulate, nanoechinate, reticulum cristatum (SEM).

Remarks: The figured specimen displays a reticulum crested by relatively long nanoechini; similar pollen morphology was reported in *Pistacia atlantica* Desf. [33].

Family Sapindaceae Juss.
Genus *Acer* L.
*Acer* sp. 1 (Figs 5e–f)

Description: Pollen, monad, shape prolate, lobate in polar view, pollen size medium, diameter 30–40 μm (LM); tricolpate; sculpturing striate (LM, SEM).

Remarks: Striate exine sculpturing is the most common type in *Acer* [34].

*Acer* sp. 2 (Figs 5g–h)

Description: Pollen, monad, shape prolate, elliptic in equatorial view, lobate in polar view, pollen size medium, diameter 30–40 μm (LM), polar axis 35–45; tricolpate; sculpturing scabrate (LM), rugulate to striatoreticulate, fossulate, perforate (SEM).

Remarks: This type of exine sculpture is only found in the infrageneric sections *Rubra* and *Negundo* [34].

Order Malvales Juss. ex Bercht. & J.Presl
Family Malvaceae Juss.
Genus *Craigia* W.W.Sm. & W.E.Evans
*Craigia* sp. (Figs 5i–j)

Description: Pollen, monad, shape oblate, circular to convex triangular in polar view, pollen size medium, diameter 25–35 μm (LM); tricolporate, colpi short, endo aperture circular to elliptic; sculpturing microreticulate, heterobrochate (LM, SEM).

Remarks: The figured specimen corresponds by exine sculpturing and aperture configuration (horseshoe-shaped thickening in aperture area in polar view) to *Craigia* [35].

Order Caryophyllales Juss. ex Bercht. & J.Presl
Family Amaranthaceae Juss.
Amaranthaceae gen. indet. 1 (Fig. 5k)

Description: Pollen, monad, shape spheroidal, outline circular, pollen size small, diameter 18–22 μm (LM); pantoporate, pori sunken, pori diameter 1–2 µm, 34–42 pori; sculpturing psilate (LM), perforate, nanoechinate (SEM).

Amaranthaceae gen. indet. 2 (Fig. 5l)

Description: Pollen, monad, shape spheroidal, outline circular, pollen size small, diameter 20–25 μm (LM); pantoporate, pori sunken, pori diameter 2–4 µm, 26–32 pori; sculpturing psilate (LM), perforate, nanoechinate (SEM).

Remarks: Amaranthaceae gen. indet. 1 differs by smaller size and pores from Amaranthaceae gen. indet. 2. [14].

Family Caryophyllaceae Juss.
Caryophyllaceae gen. indet. (Figs 5m–n)

Description: Pollen, monad, shape spheroidal, outline circular, pollen size medium, diameter 35–50 μm (LM); pantoporate, pori sunken, pori diameter 5–7 µm, 12 pori, porus membrane with 8–11 microechini; sculpturing psilate (LM), perforate, nanoechinate (SEM).

Remarks: The figured specimen is poorly preserved, but shares morphological similarities (absence of distinct annulus, evenly distributed nanoechini and perforations, pore membrane covered by microechini) with the Cerastium fontanum type [36].

Order Cornales Link
Family Nyssaceae Dumortier
Genus *Nyssa* L.
*Nyssa* sp. (Figs 5o–p)

Description: Pollen, monad, shape spheroidal to subprolate, outline circular to convex triangular in polar view, circular in equatorial view, pollen size medium, polar axis 35–50 μm long (LM); tricolporate, endoporus circular; sculpturing scabrate (LM), rugulate, fossulate, perforate (SEM), perforations absent in colpus area.

Remarks: Pollen with similar smooth colpus area have been reported in extant *Nyssa sylvatica* Marshall and *N. sinensis* Oliv. [37].

Order Lamiales Bromhead
Family Oleaceae Hoffmanns. & Link
Genus *Fraxinus* L.
*Fraxinus* sp. (Figs 5q–r)

Description: Pollen, monad, shape prolate, circular in equatorial view, pollen size medium, equatorial diameter 20–30 µm (LM), polar axis 25–40 μm long (LM); tricolpate; sculpturing reticulate, heterobrochate (LM, SEM), reticulum crested by blunt nano echini and perpendicular ridges (SEM, Fig. 5r).

Remarks: The figured specimen corresponds (size, reticulum morphology, aperture) to the pollen of this genus [38, 39]. Morphologically similar pollen is produced by extant *Fraxinus excelsior* [40]. Fossil seeds of *Fraxinus* have been reported from the Vegora mine [41].

Genus *Olea* L.
*Olea* sp. (Figs 5s–t)

Description: Pollen, monad, shape spheroidal, circular in equatorial and polar view, pollen size small, equatorial diameter 18–25 µm (LM), polar axis 20–25 μm long (LM); tricolporate; sculpturing reticulate, heterobrochate (LM, SEM), reticulum crested by blunt nano echini (SEM, Fig. 5t).

Remarks: The figured specimen corresponds (reticulum morphology, aperture) to pollen of this genus [38, 39].

Order Asterales Link
Family Asteraceae Bercht. & Presl
Subfamily Asteroideae Lindl. in Loudon
Asteroideae gen. indet. sp. 1 (Fig. 5v)

Description: Pollen, monad, shape spheroidal, circular in equatorial, lobate in polar view, pollen size medium, diameter 25–30 µm (LM), polar axis 25–30 μm long (LM); tricolporate; sculpturing echinate, (LM), perforations extend to the upper third of echini, perforations circular (SEM).

Remarks: The poor preservation of Asteroideae gen. indet. sp. 1 and 2 prevents the determination of the aperture configuration, which is vital for the assignment to genus level.

Asteroideae gen. indet. sp. 2 (Fig. 5w)

Description: Pollen, monad, shape spheroidal, circular in equatorial, lobate in polar view, pollen size medium, diameter 30–40 µm (LM), polar axis 30–40 μm long (LM); tricolporate; sculpturing echinate, (LM), perforations extend to the upper third of echini, perforations of irregular shape (SEM).

Subfamily Cichorioideae Chevall.
Cichorioideae gen. indet. sp. (Fig. 5u)

Description: Pollen, monad, shape spheroidal, outline polygonal, pollen size medium, diameter 25–35 µm (LM), tricolporate; sculpturing echinate, lophate, (LM, SEM).

Remarks: Lophate pollen is characteristic of Cichorioideae [42]. The poor preservation prevents assignment to genus level.

Order Dipsacales Juss. ex Bercht. & Presl
Family Caprifoliaceae Juss.
Genus Succisa Moench
*Succisa* sp. (Figs 5x–z)

Description: Pollen, monad, shape oblate to subspheroidal, outline circular to weakly lobate in polar view, pollen size large, diameter 60–75 µm (LM), tricolpate, colpus membrane echinate, echini 1.5–2.5 µm long; sculpturing echinate, microechinate, echini irregularly distributed over exine surface (LM, SEM), indistinct perforate (SEM).

Remarks: The morphology of the figured specimen corresponds (aperture without halo, echinate colpus membrane, irregularly distributed echini and microechini) to the *Succisa pratense* type of Clarke and Jones [43].

Order Apiales Nakai
Family Apiaceae Lindl.
Apiaceae sp.1 (Figs 5aa–bb)

Description: Pollen, monad, shape prolate, outline elliptic in equatorial view, pollen size small to medium, equatorial diameter 9–12 µm (LM), polar axis 22–30 µm (LM), tricolporate, colpus length 3/4 to 5/6 of polar axis, endoporus lalongate elliptic; sculpturing scabrate (LM), microrugulate, fossulate, perforate (SEM).

Remarks: Morphological similarities (colpus length, endoporus shape, exine sculpture) are with the *Sium latifolium* type of Punt [44].

Apiaceae sp. 2 (Figs 5cc–dd)

Description: Pollen, monad, shape prolate, outline bone-shaped in equatorial view, pollen size small, equatorial diameter 6–8 µm (LM), polar axis 16–20 µm (LM), tricolporate, colpus length 1/3 to 1/2 of polar axis, endoporus lalongate elliptic; sculpturing scabrate (LM), perforate (SEM).

Remarks: Morphological similarities (colpus length, endoporus shape, exine sculpture) are with the *Trinia glauca* type and the *Sison amomum* type of Punt [44].

Incertae sedis

Monocotyledonae indet. (Figs 4d–e)

Description: Pollen, monad, shape oblate, outline elliptic in polar view, monad size small, diameter 18–25 μm (LM); sulcate; sculpturing reticulate (LM, SEM).

Remarks: The figured specimen is not complete; therefore unambiguous determination is not possible. Sulcate, reticulate pollen is commonly present in e.g. Arecaceae [45] or Liliaceae [46].

Dicotyledonae fam. et gen. indet. (Figs 4ee–ff)

Description: Pollen, monad, shape prolate, outline elliptic in equatorial view, pollen size small to medium, diameter 20–30 µm (LM), tricolporate; sculpturing scabrate (LM), microreticulate, reticulum with perpendicular ridges (SEM).
