## Supplemental Material 5 for "Messinian vegetation and climate of the intermontane Florina-Ptolemais-Servia Basin, NW Greece inferred from palaeobotanical data: How well do plant fossils reflect past environments?": Physg3arcAZ_GRIDMet3arAZ.pdf (from another version of this preprint: version 1, 2019-11-25)

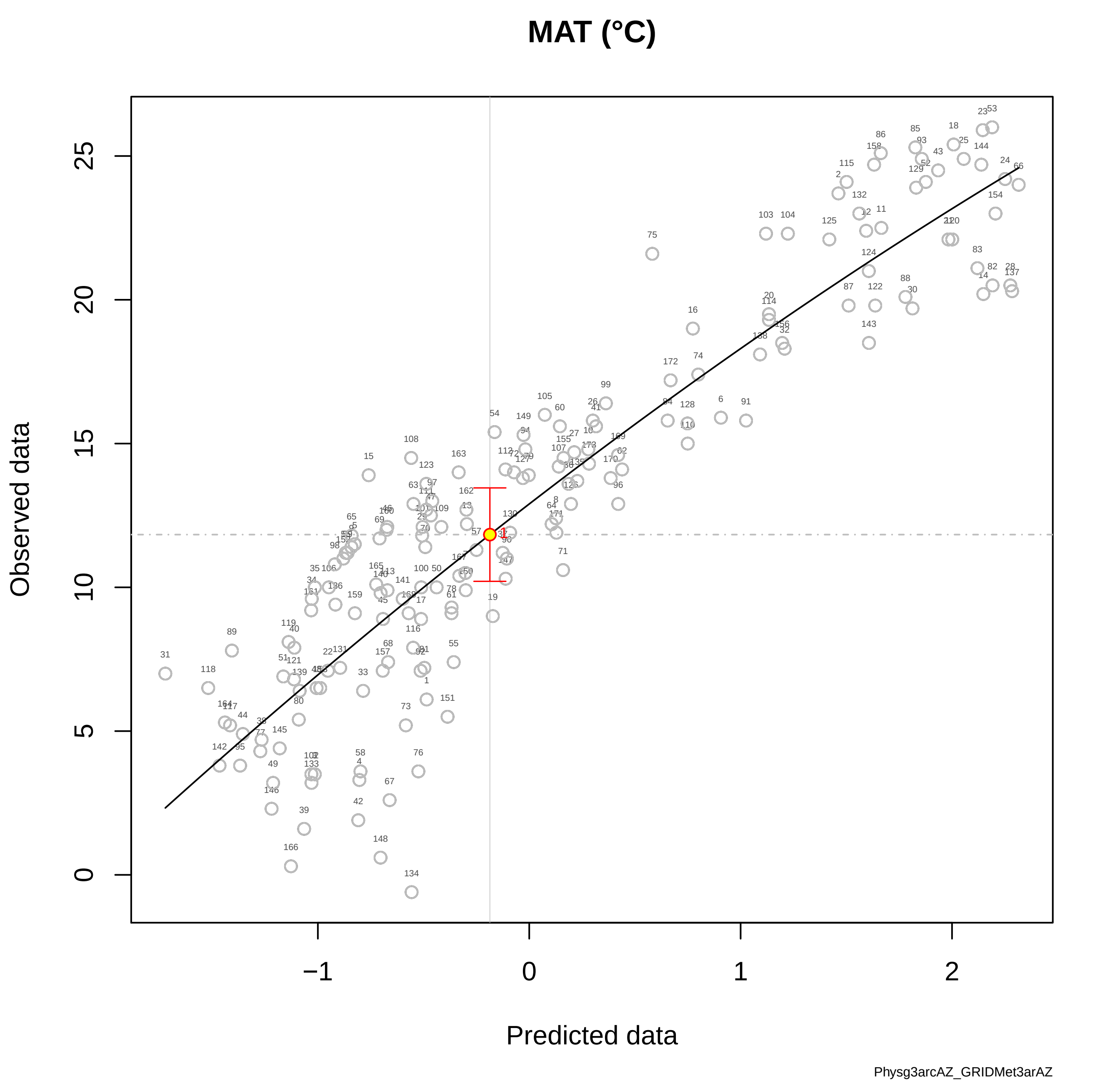
MAT (°C)
Observed data
Predicted data
0
5
10
15
20
25
−1
0
1
2
Physg3arcAZ_GRIDMet3arAZ

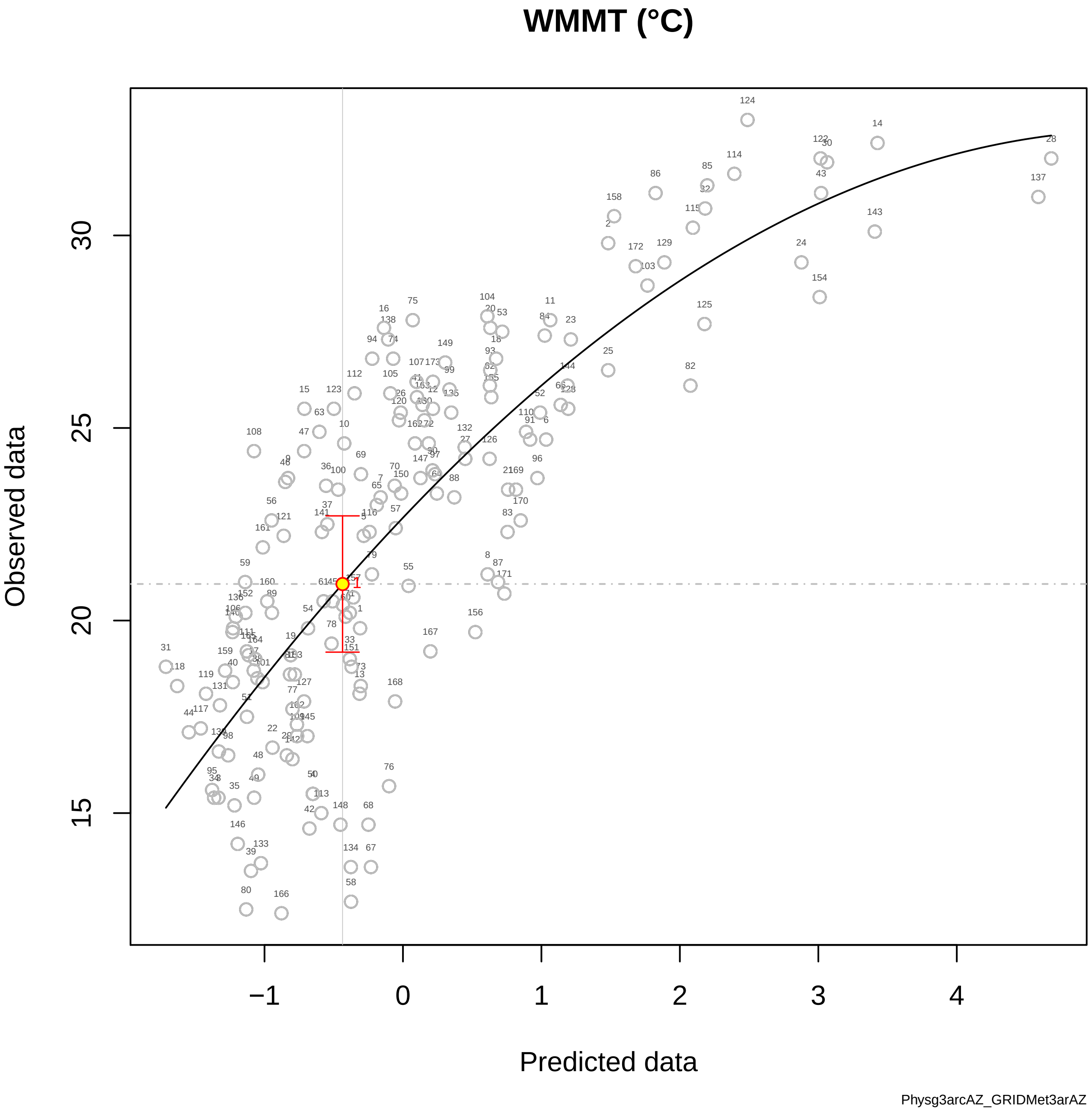
WMMT (°C)
Observed data
Predicted data
15
20
25
30
−1
0
1
2
3
4
Physg3arcAZ_GRIDMet3arAZ

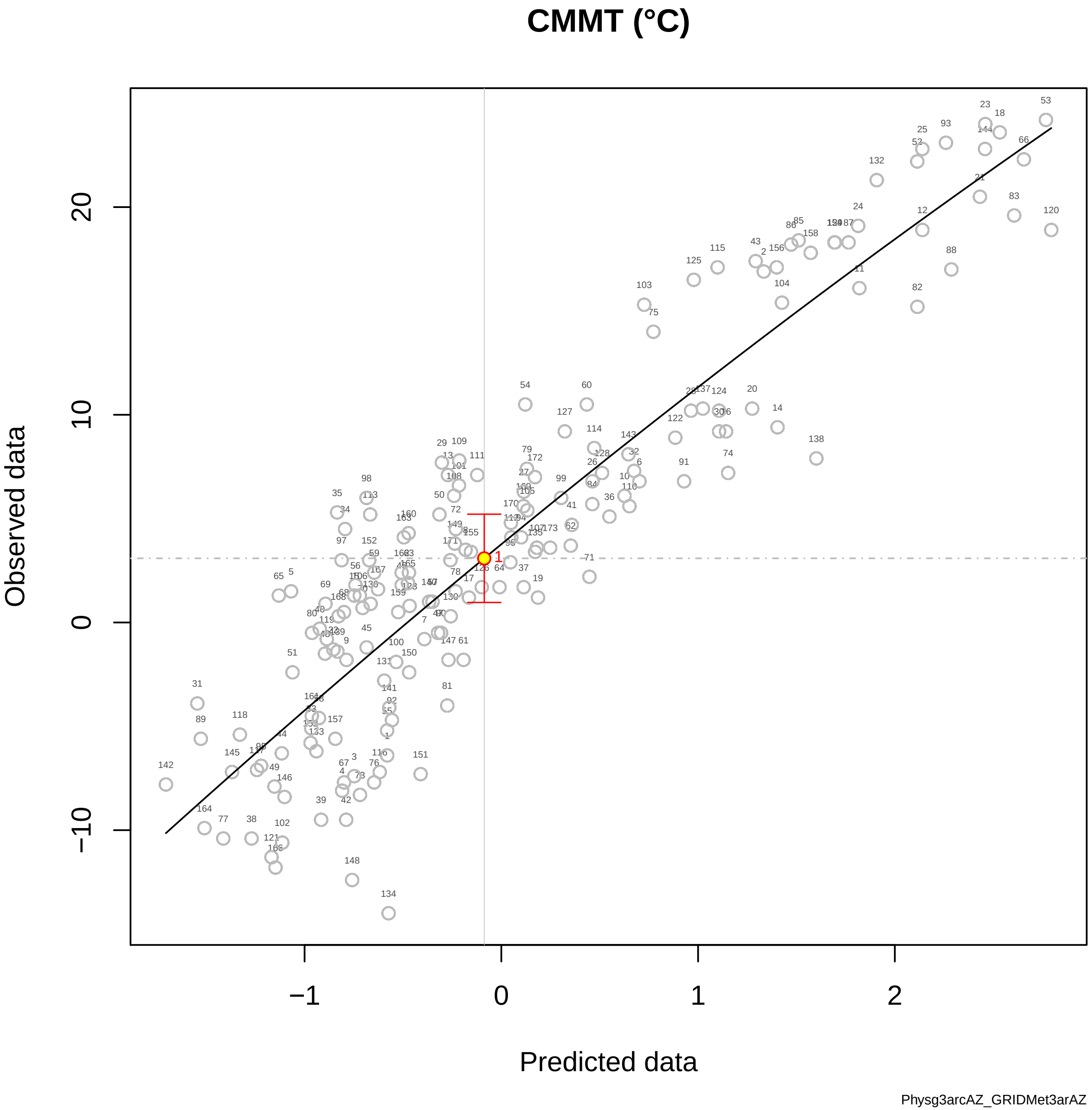
CMMT (°C)
Observed data
Predicted data
20
10
0
−10
−1
0
1
2
Physg3arcAZ_GRIDMet3arAZ

### GROWSEAS (months)

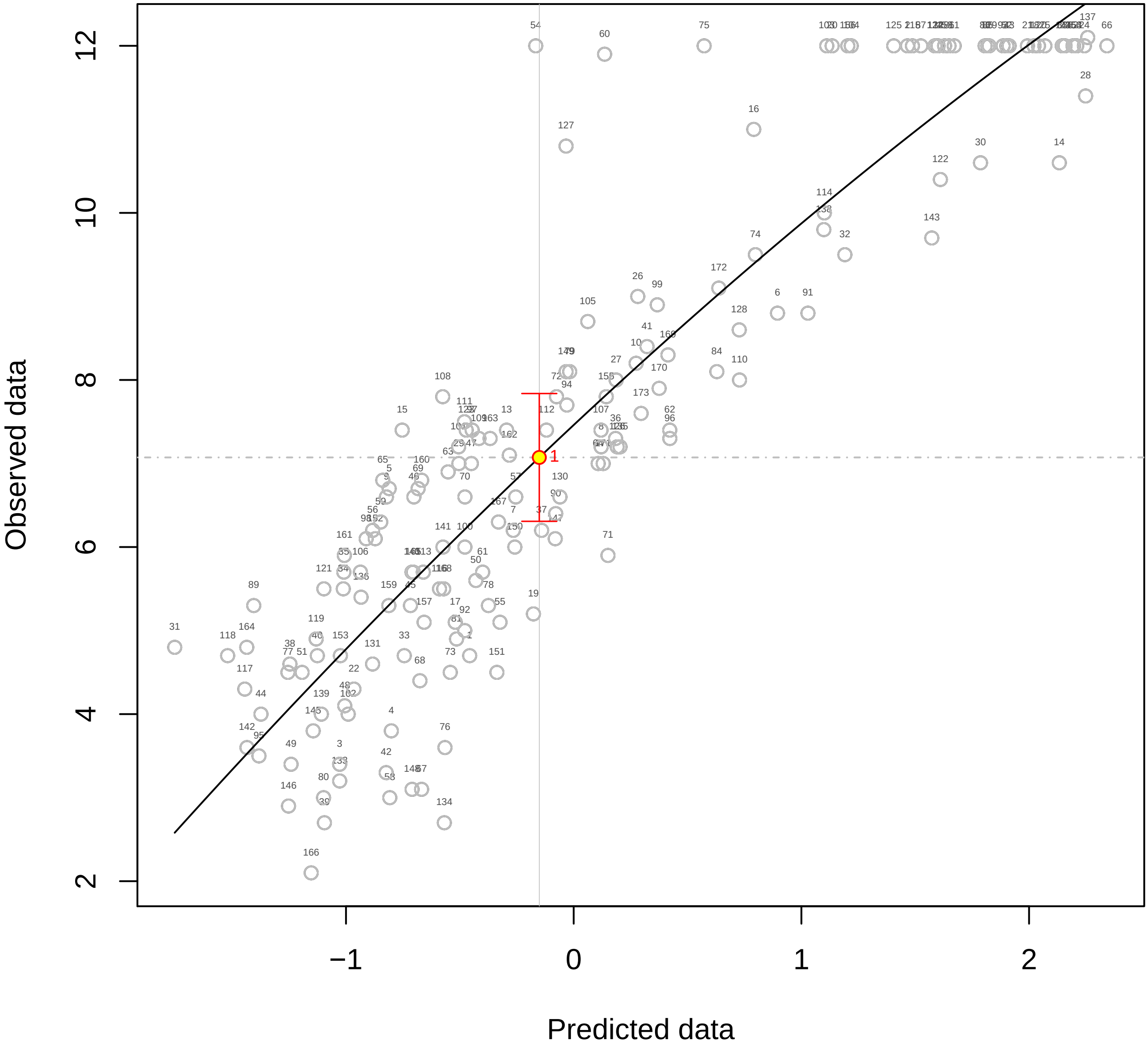


Physg3arcAZ_GRIDMet3arAZ

GSP (cm)

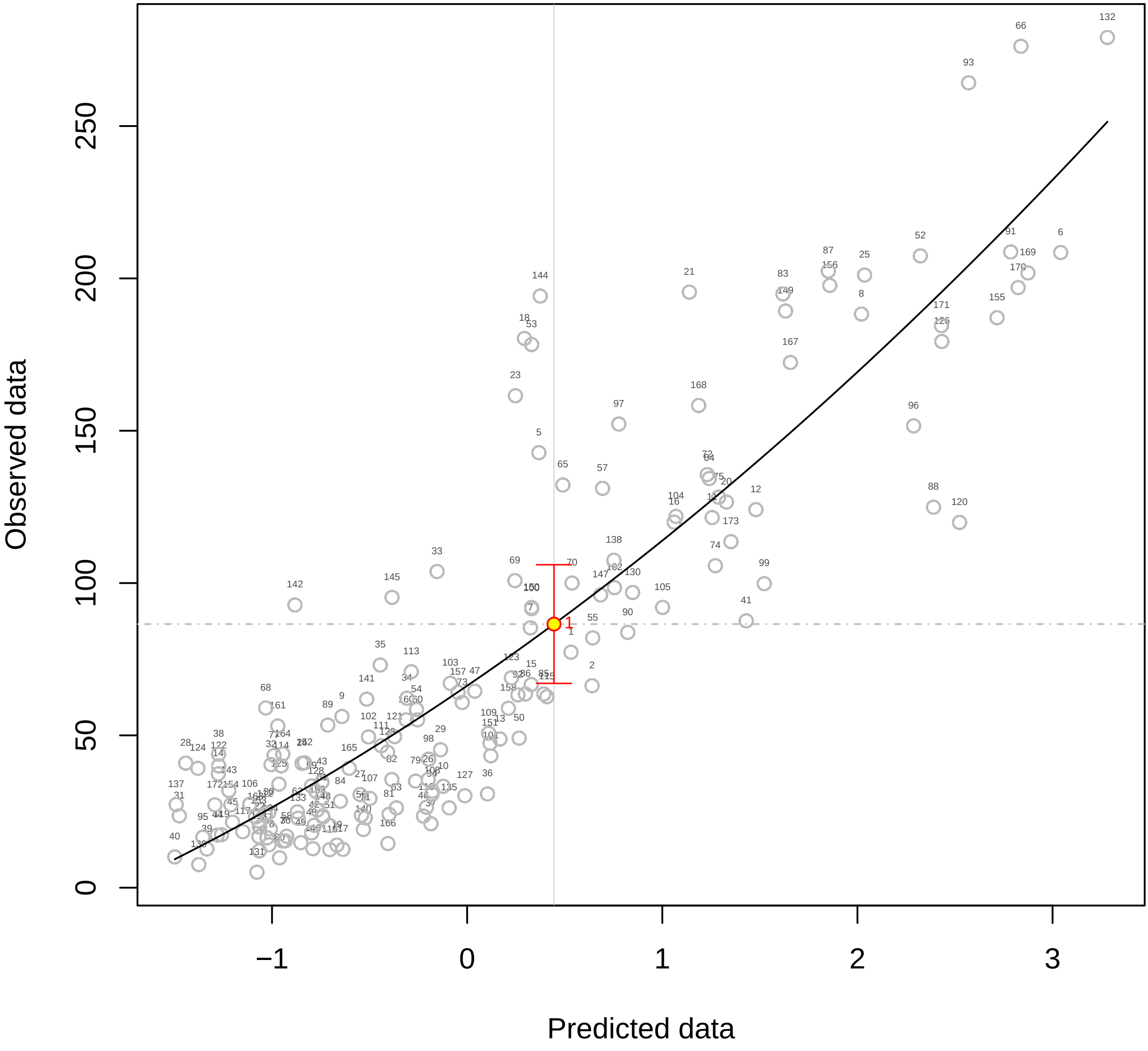


Physg3arcAZ_GRIDMet3arAZ

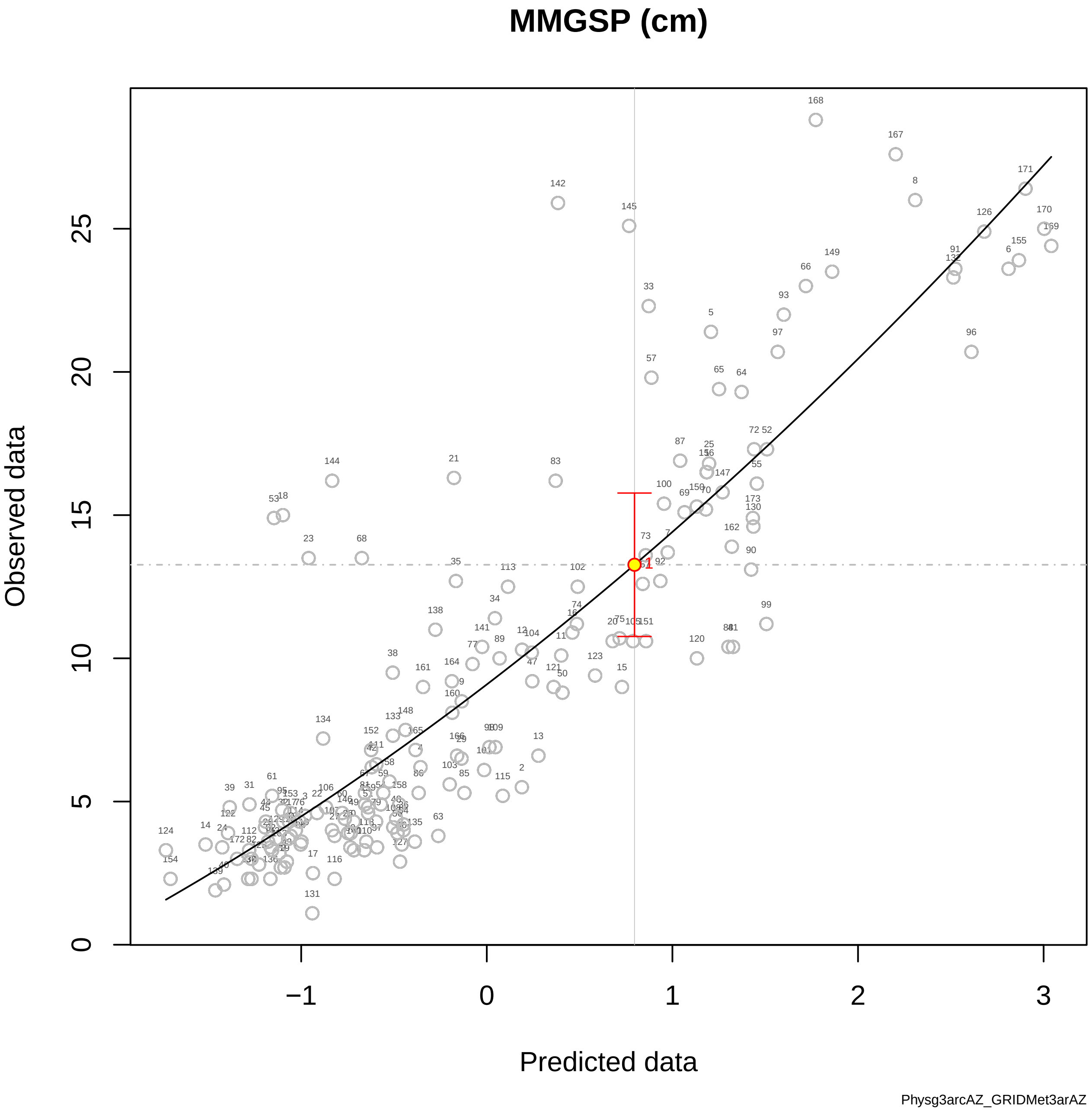
MMGSP (cm)
Observed data
Predicted data
0
5
10
15
20
25
−1
0
1
2
3
Physg3arcAZ_GRIDMet3arAZ

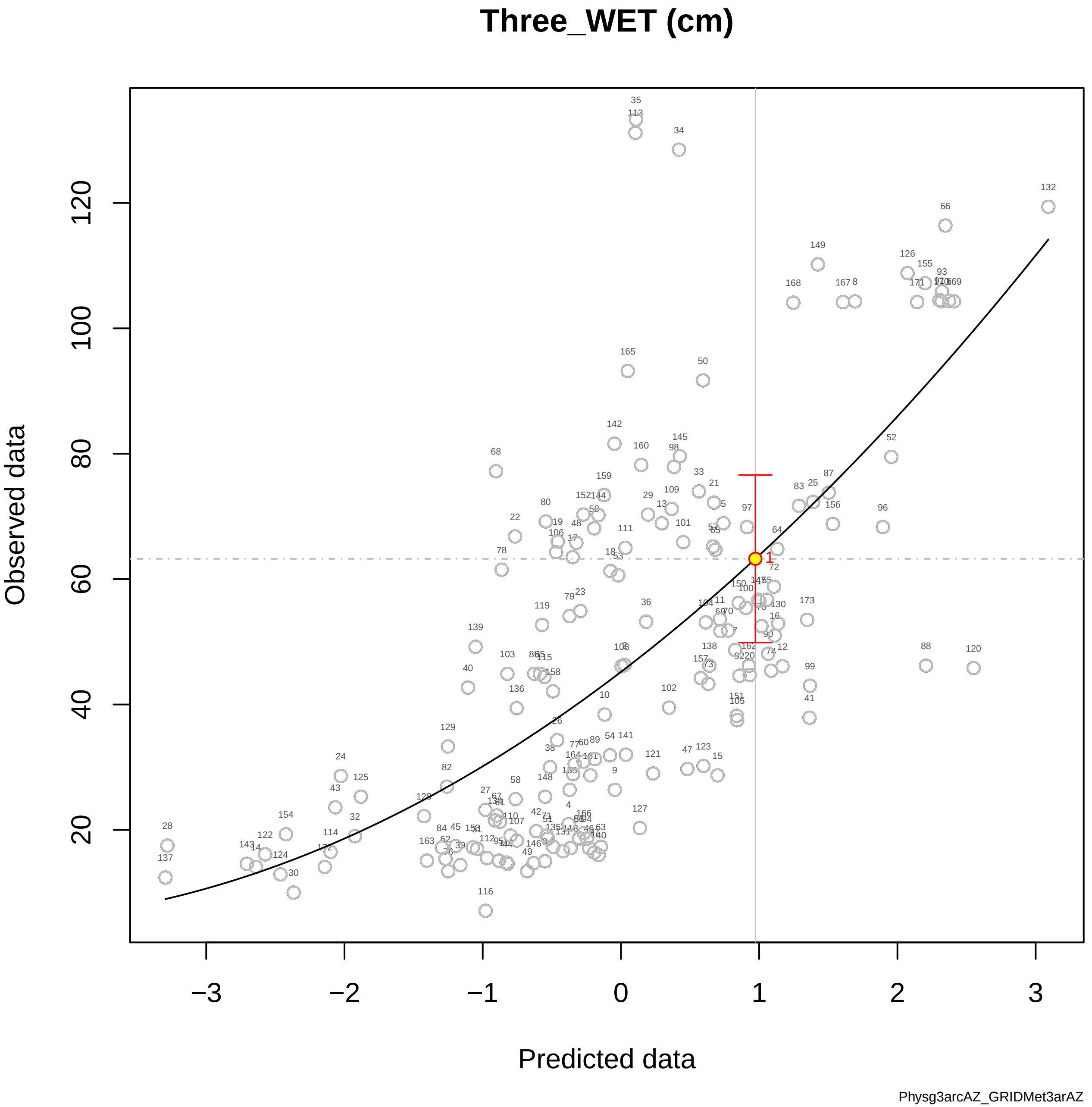
Three_WET (cm)
Observed data
Predicted data
Physg3arcAZ_GRIDMet3arAZ

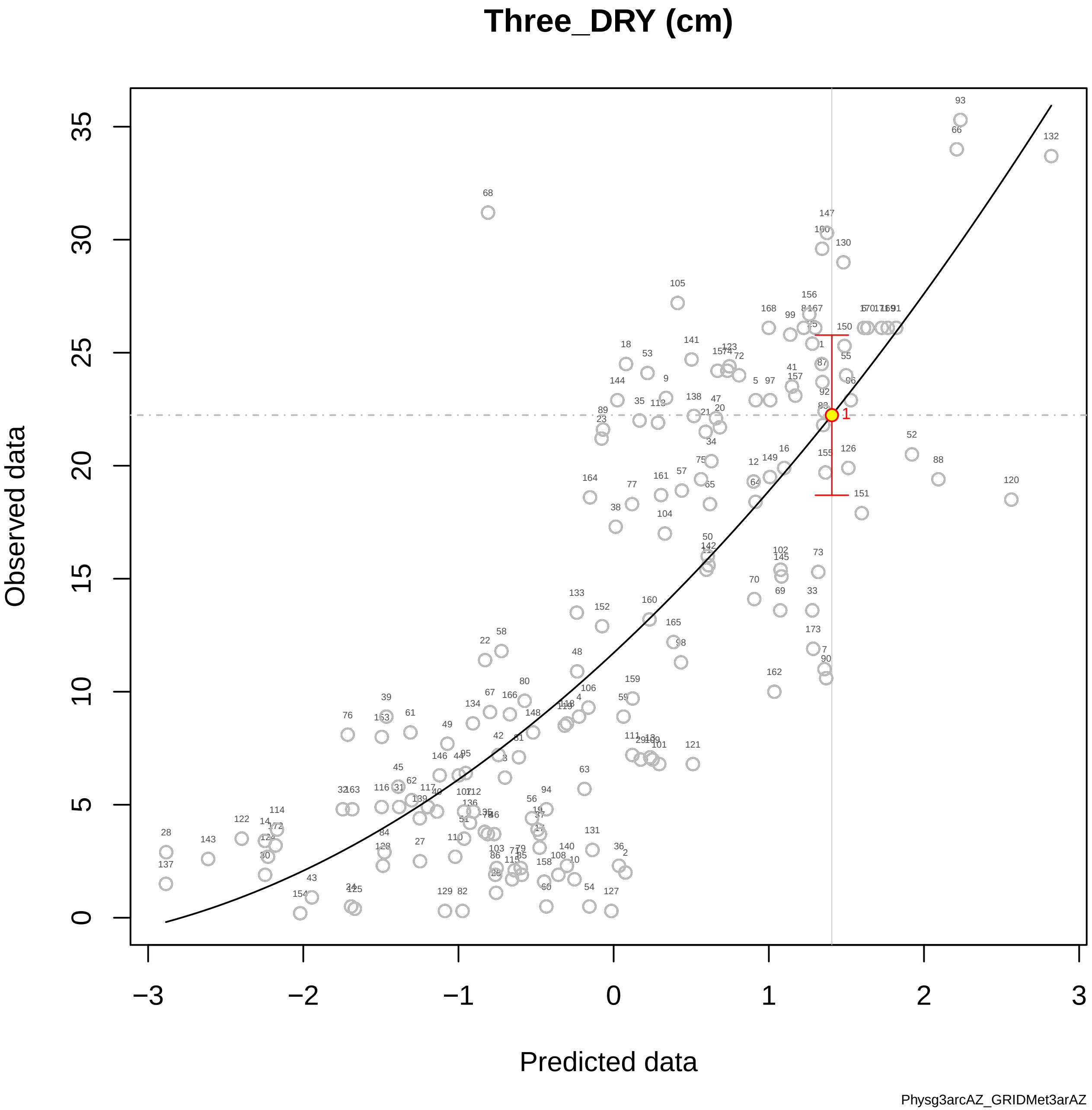
Three_DRY (cm)
Observed data
Predicted data
Physg3arcAZ_GRIDMet3arAZ

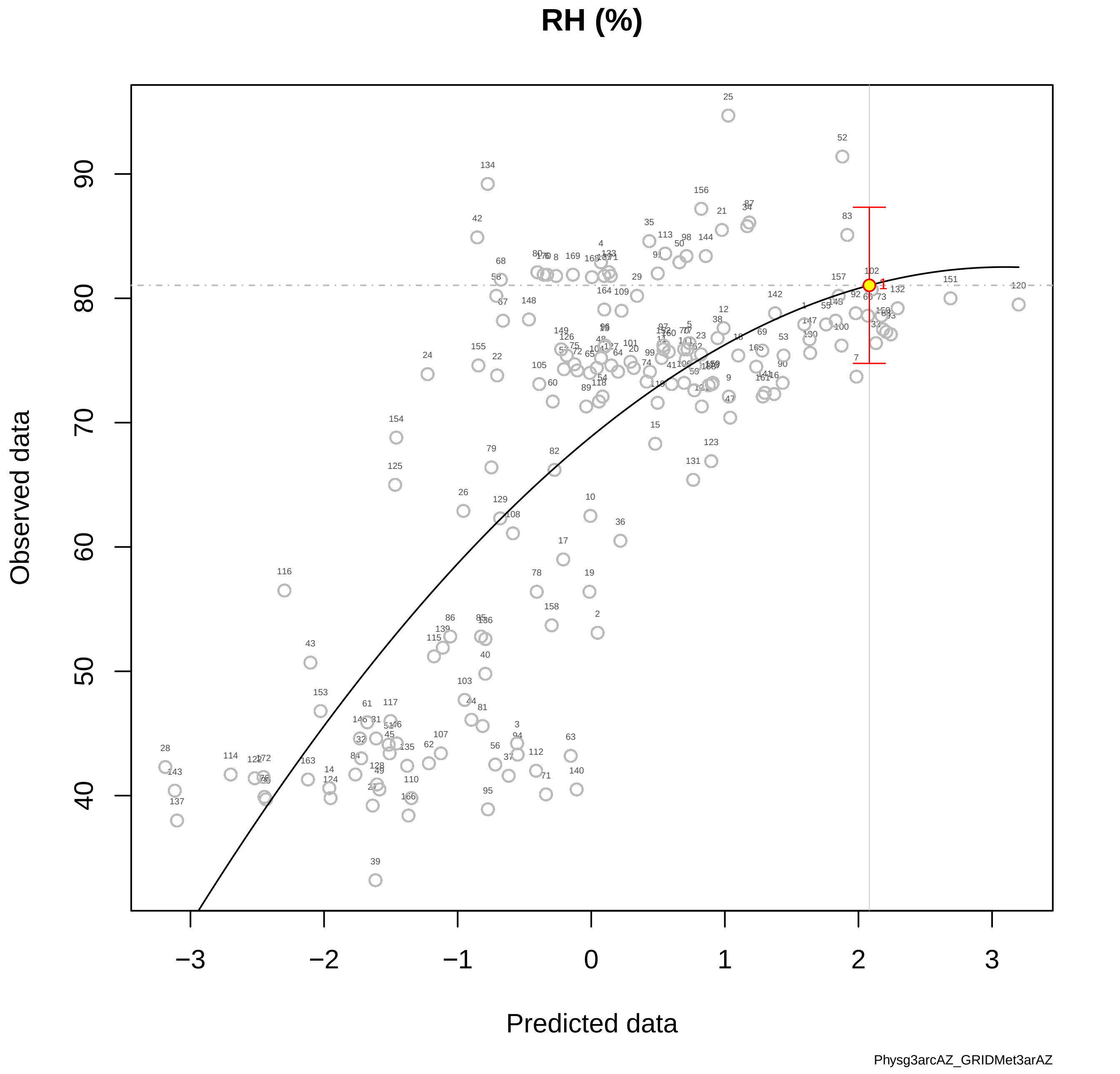
RH (%)
Observed data
Predicted data
90
80
70
60
50
40
−3
−2
−1
0
1
2
3
Physg3arcAZ_GRIDMet3arAZ

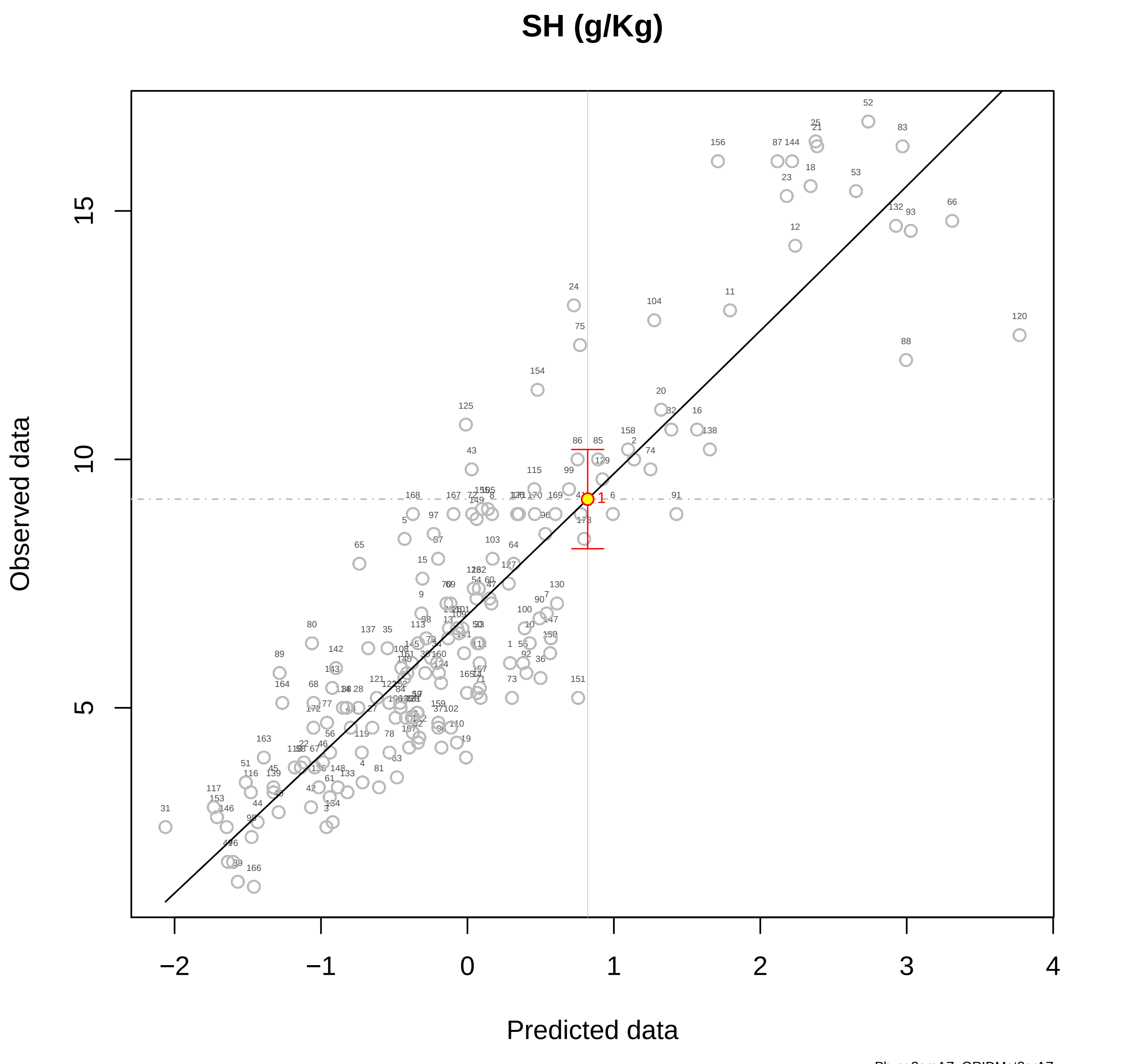

Physg3arcAZ_GRIDMet3arAZ

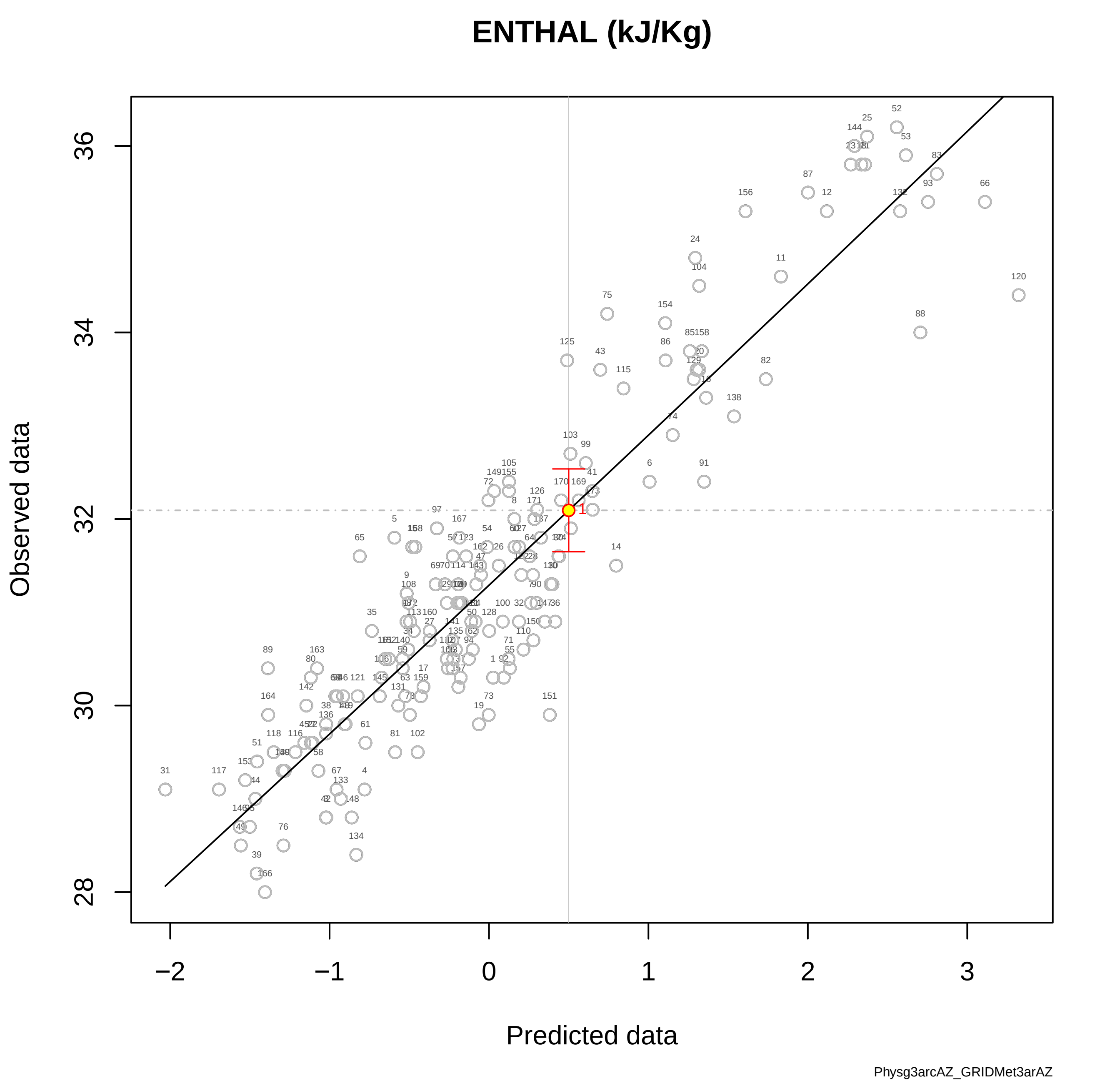
ENTHAL (kJ/Kg)
Observed data
Predicted data
36
34
32
30
28
−2
−1
0
1
2
3
Physg3arcAZ_GRIDMet3arAZ
