## Supplemental Material 5 for "Messinian vegetation and climate of the intermontane Florina-Ptolemais-Servia Basin, NW Greece inferred from palaeobotanical data: How well do plant fossils reflect past environments?": Results.html (from another version of this preprint: version 1, 2019-11-25)

CLAMP\_Analysis\_Online

|  |
| --- |
| CLAMP\_Analysis\_Online |

---

|  |  |
| --- | --- |
| Results for submission: 38B5DDBD0A4397C0 |  |
| CCA1vsCCA2.jpg  CCA1vsCCA3.jpg  CCA2vsCCA3.jpg  CCA3d.jpg  CCAGraph.pdf  GSP.jpg  MAT.jpg  models.pdf  params\_38B5DDBD0A4397C0.txt  predict.csv  scoreres.csv  Vegora.csv | Parameter values: withlabel = true  calibration\_option = 1  image\_type = colour  file\_uploaded = Vegora.csv |
