## Supplemental Material 5 for "Messinian vegetation and climate of the intermontane Florina-Ptolemais-Servia Basin, NW Greece inferred from palaeobotanical data: How well do plant fossils reflect past environments?": CCAGraph.pdf (from another version of this preprint: version 1, 2019-11-25)

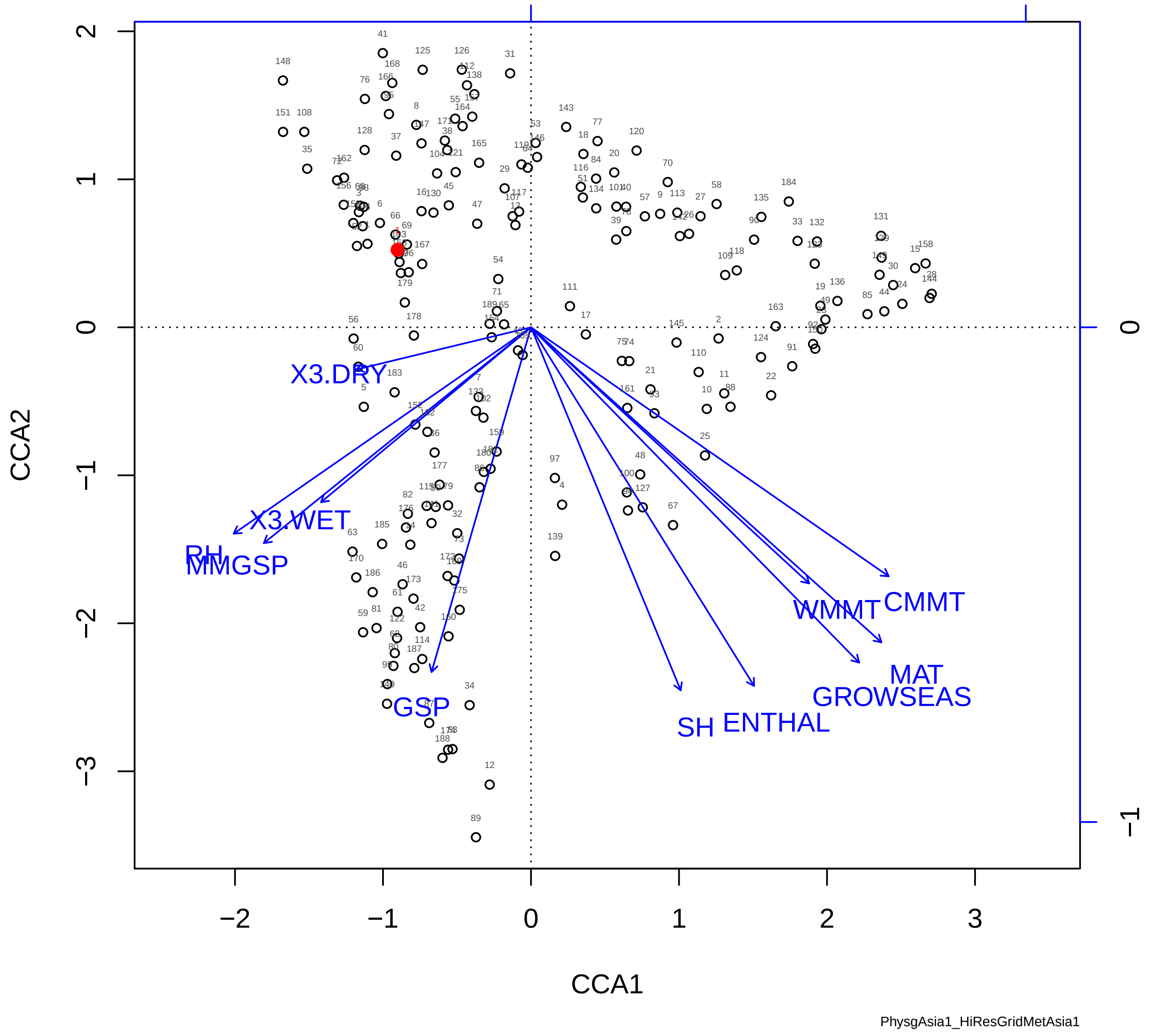

CCA1
CCA2
X3.DRY
X3.WET
RH
MMGSP
GSP
SH
ENTHAL
MAT
GROWSEAS
WMMT
CMMT
PhysgAsia1_HiResGridMetAsia1

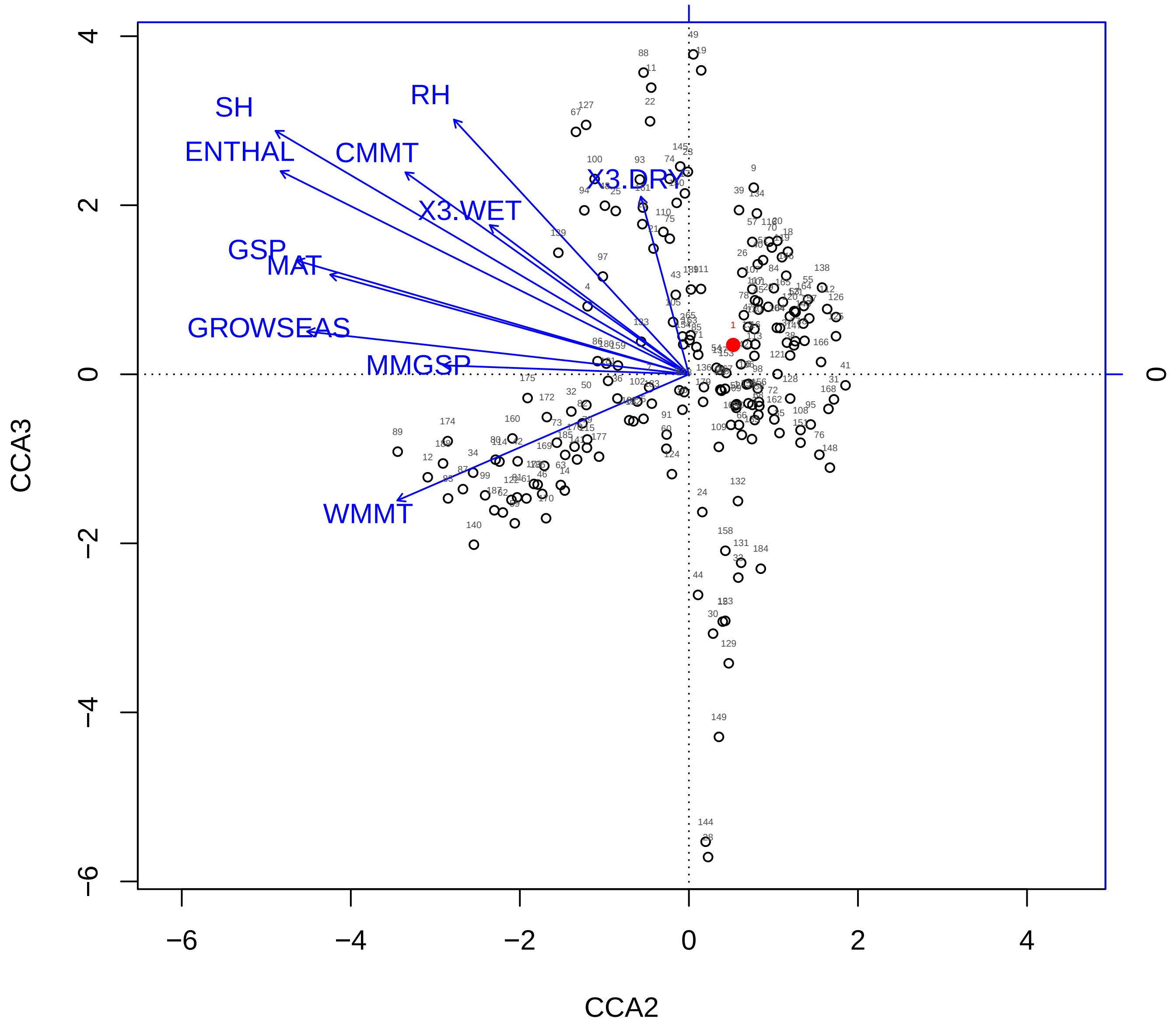

PhysgAsia1_HiResGridMetAsia1

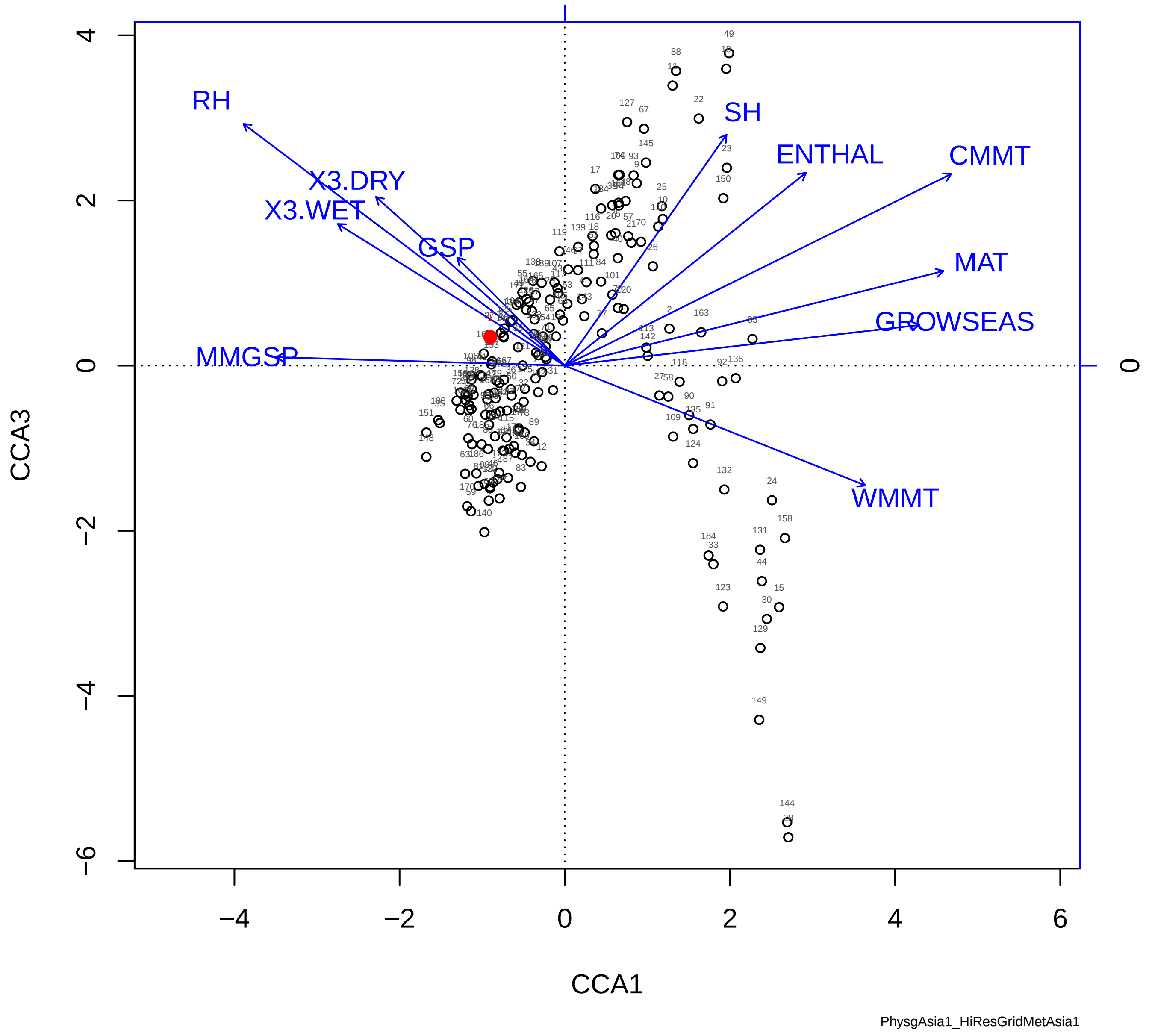

RH
X3.DRY
X3.WET
GSP
MMGSP
SH
ENTHAL
CMMT
MAT
GROWSEAS
WMMT
CCA1
CCA3
PhysgAsia1_HiResGridMetAsia1

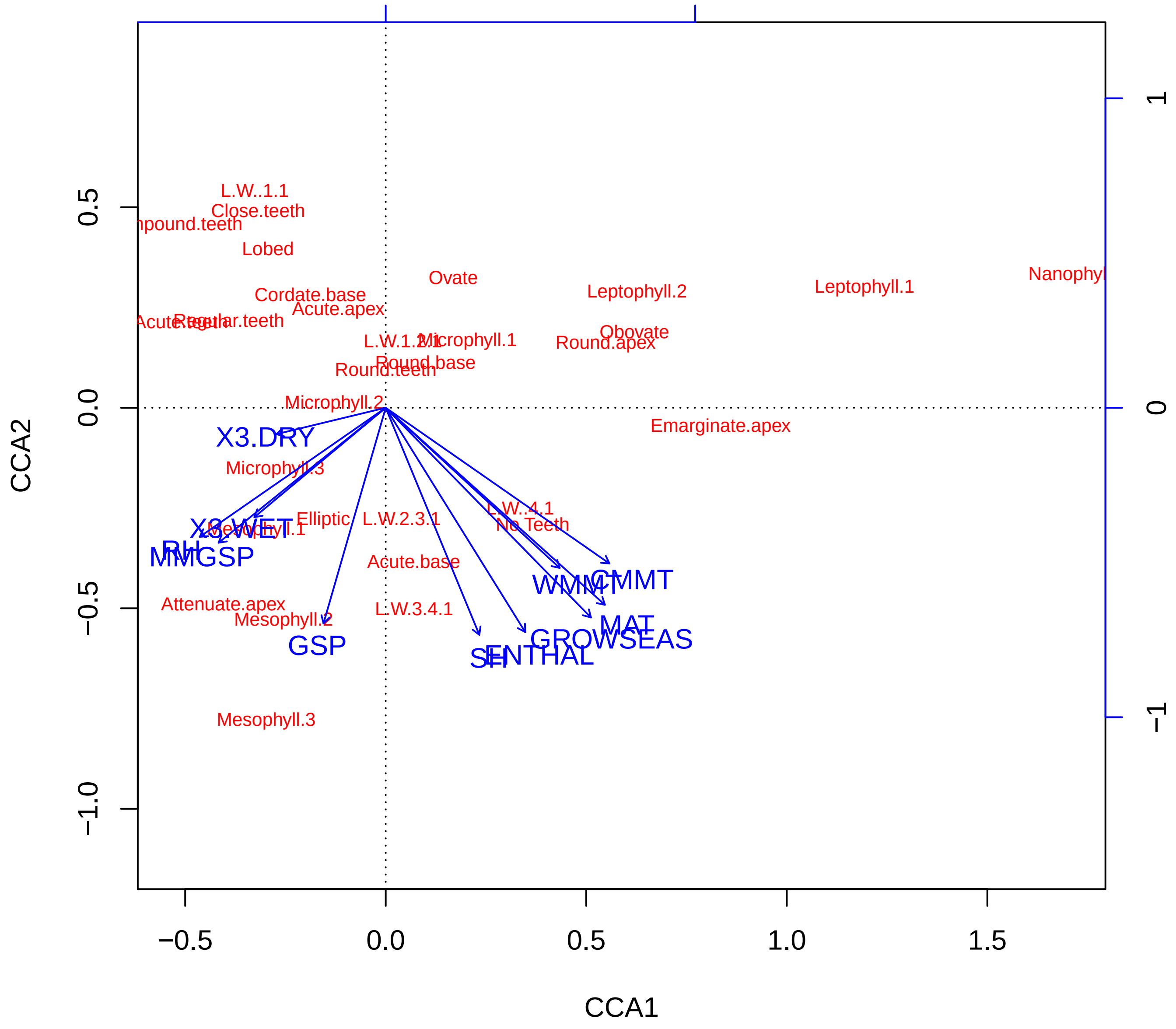

PhysgAsia1_HiResGridMetAsia1

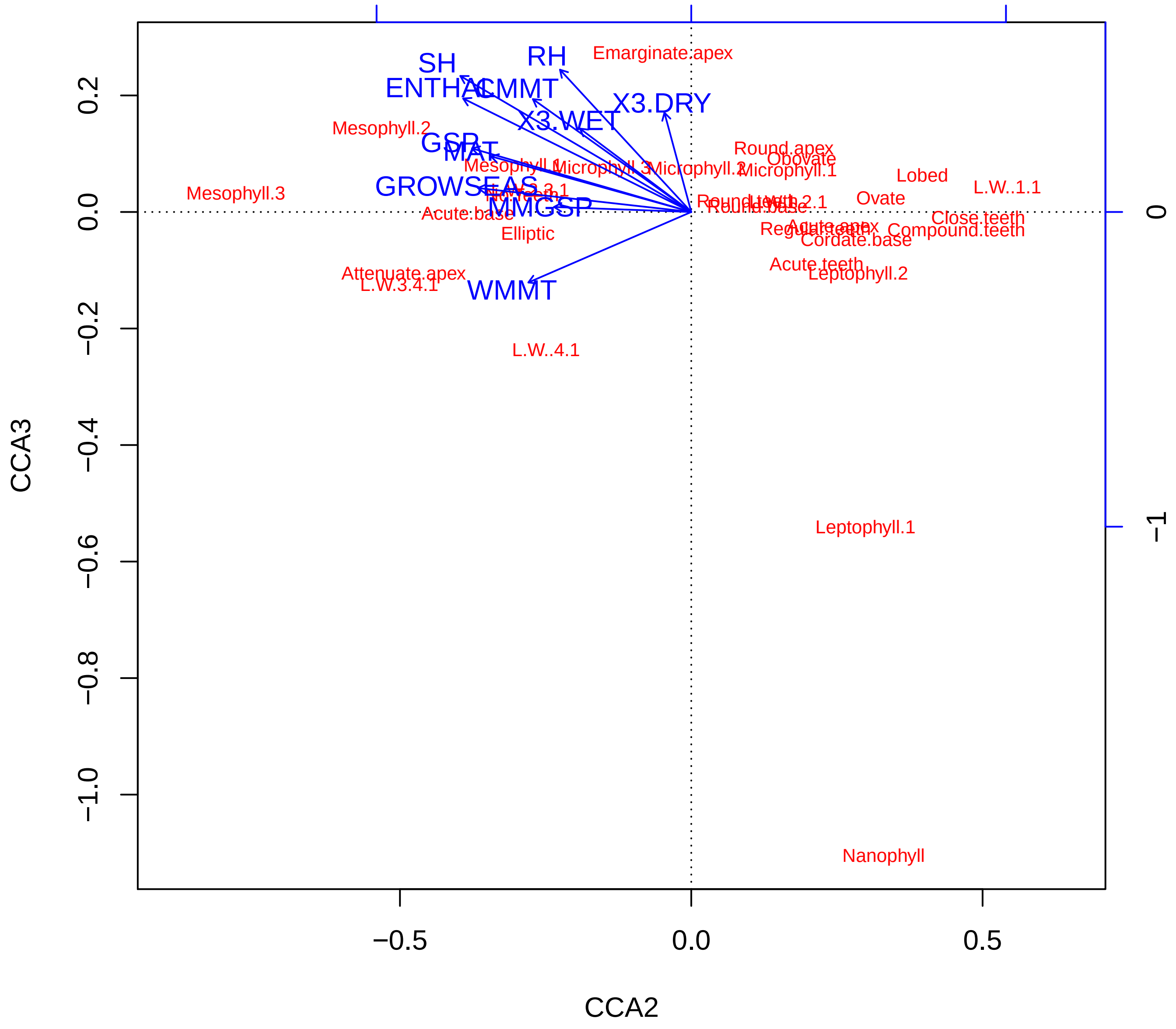

PhysgAsia1_HiResGridMetAsia1

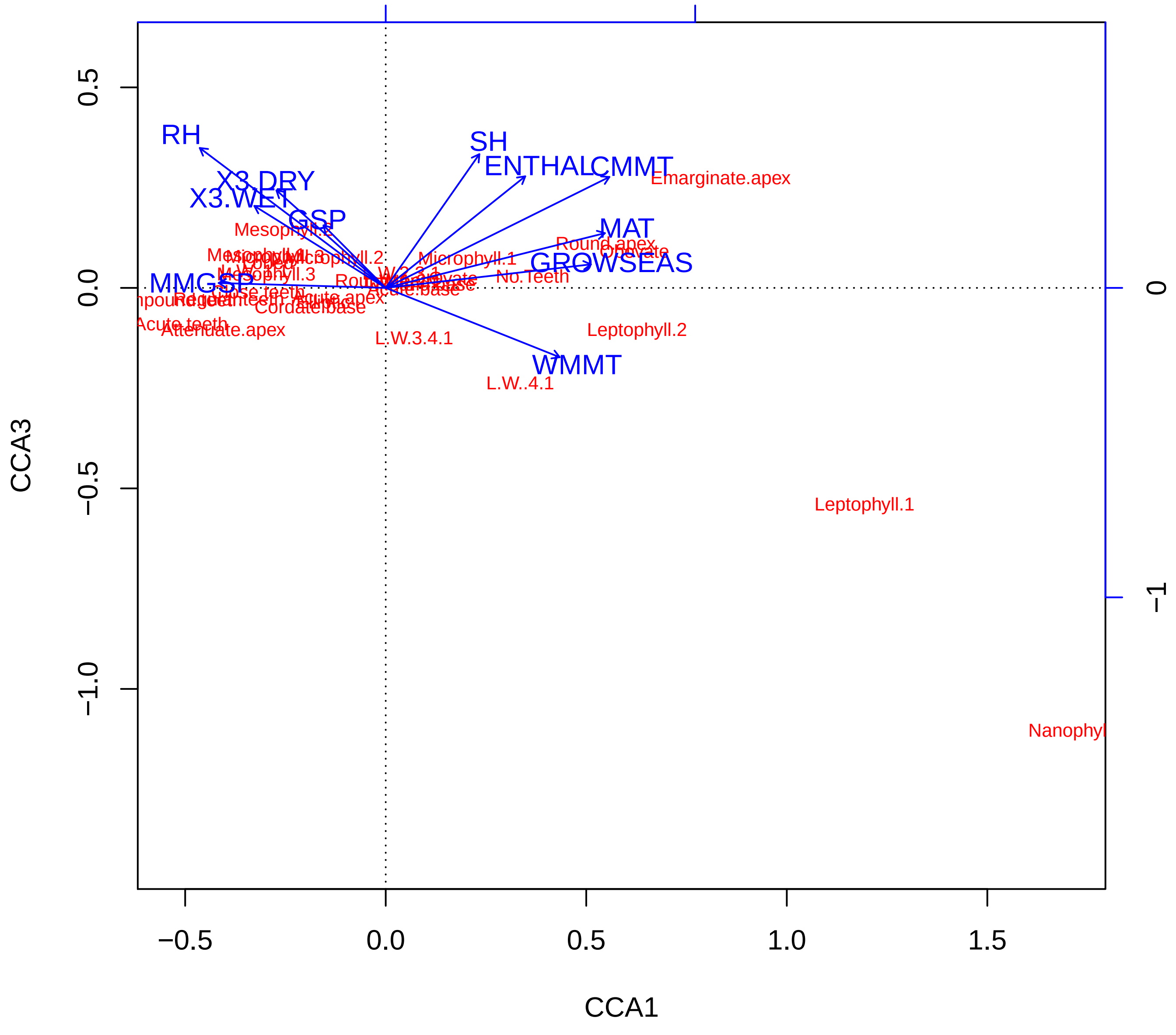


PhysgAsia1_HiResGridMetAsia1

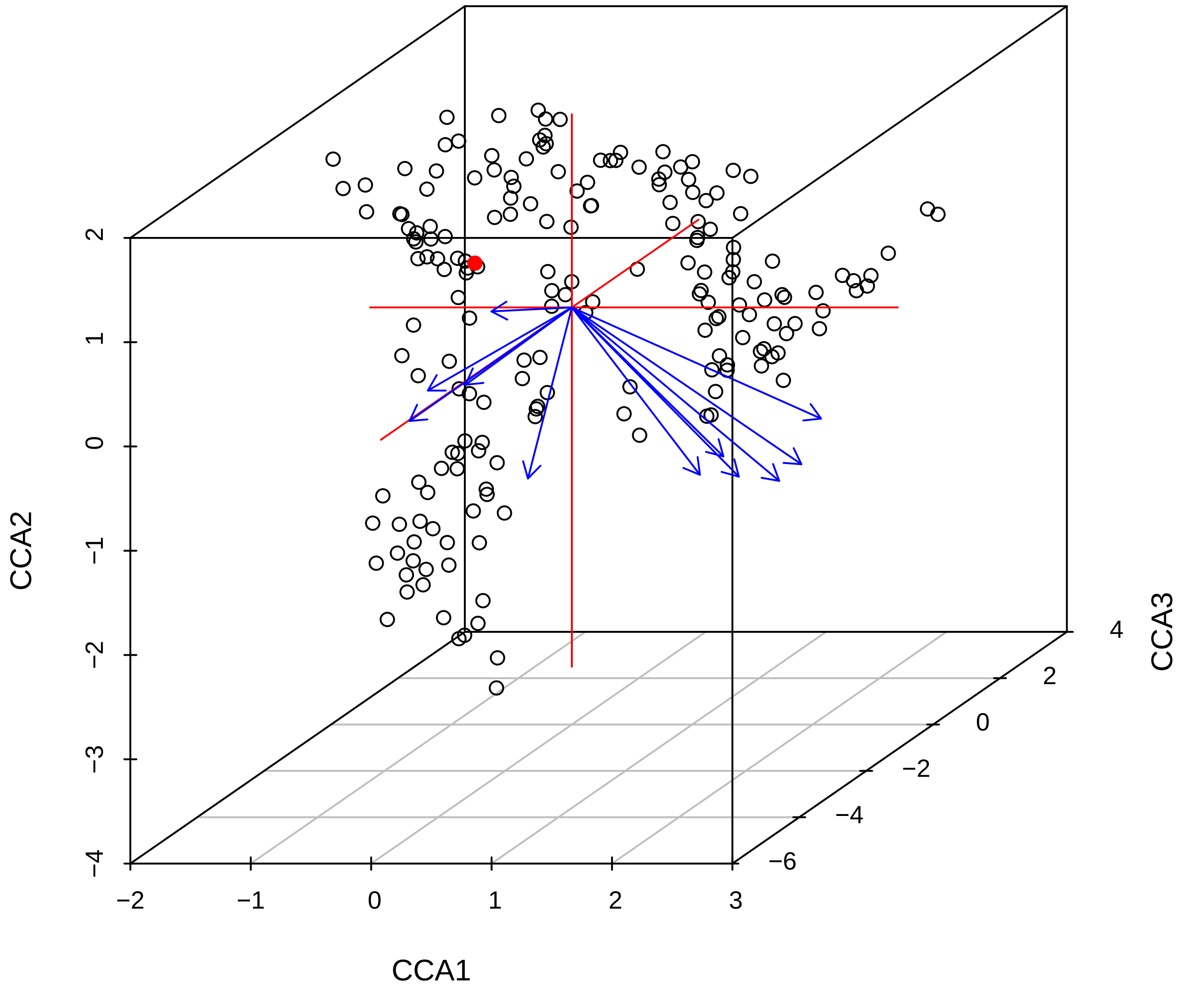


PhysgAsia1_HiResGridMetAsia1
