## Supplemental Material 5 for "Messinian vegetation and climate of the intermontane Florina-Ptolemais-Servia Basin, NW Greece inferred from palaeobotanical data: How well do plant fossils reflect past environments?": PhysgAsia1_HiResGridMetAsia1.pdf (from another version of this preprint: version 1, 2019-11-25)

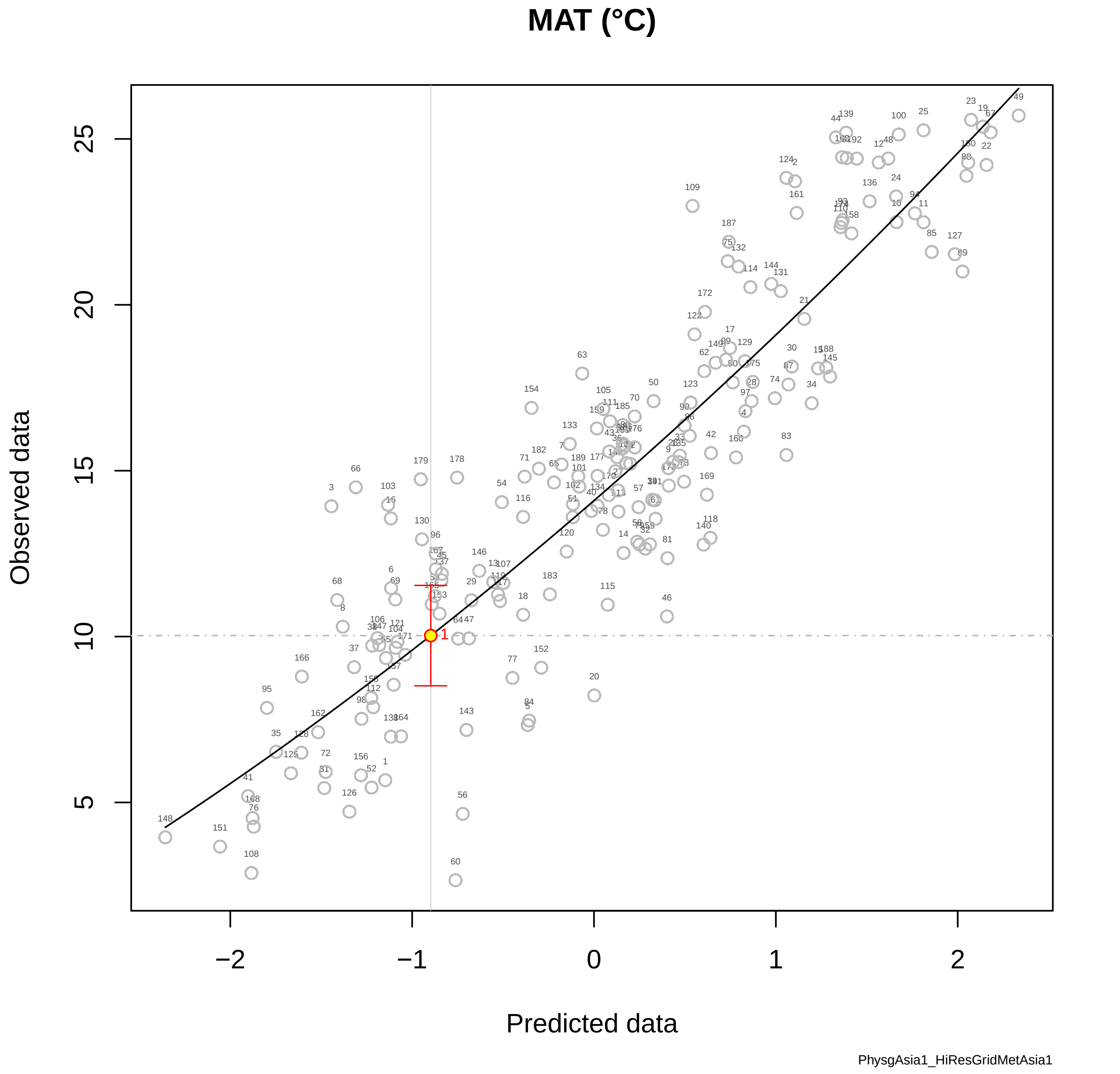
MAT (°C)
Observed data
Predicted data
PhysgAsia1_HiResGridMetAsia1

### WMMT (°C)

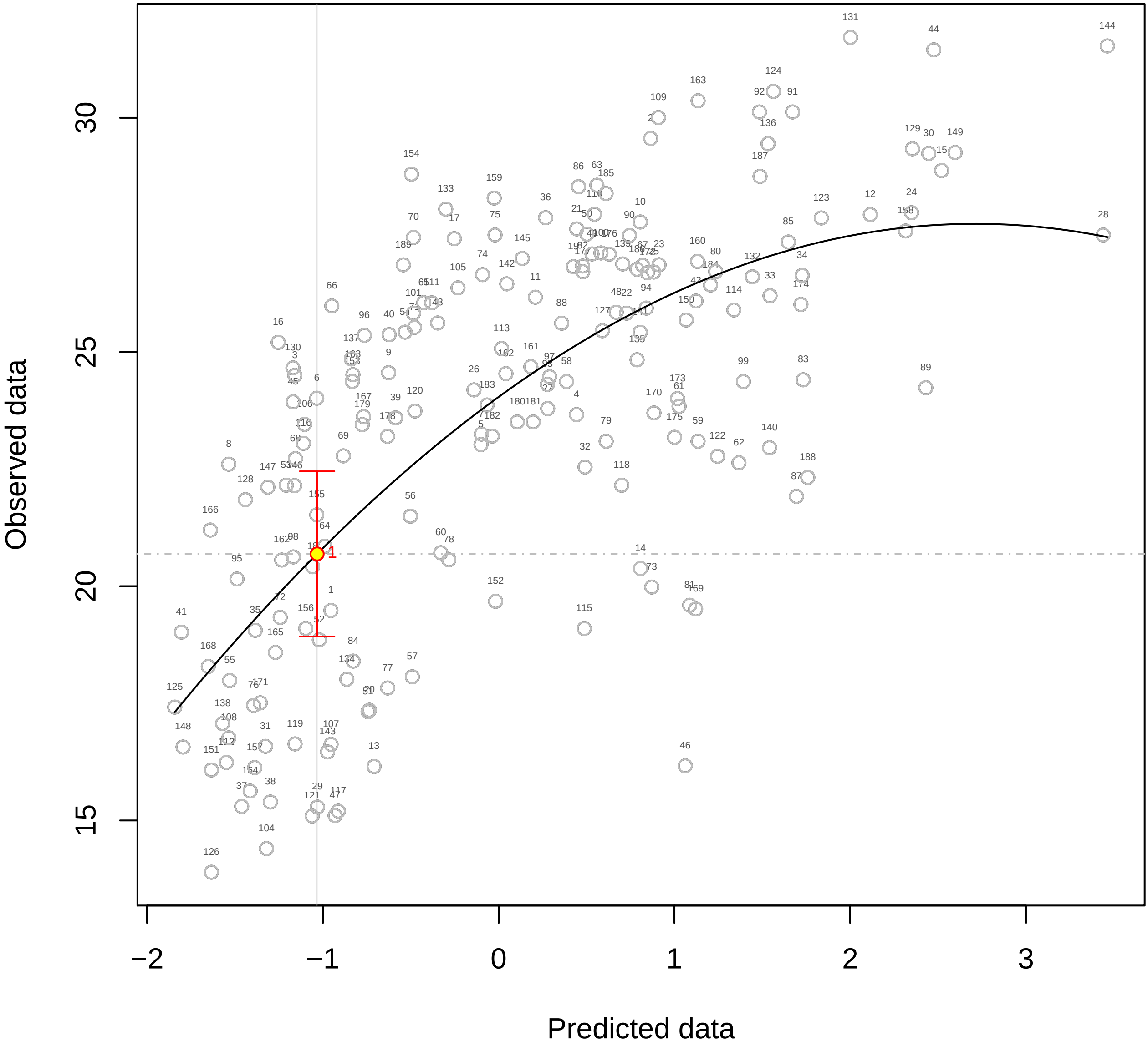


PhysgAsia1_HiResGridMetAsia1

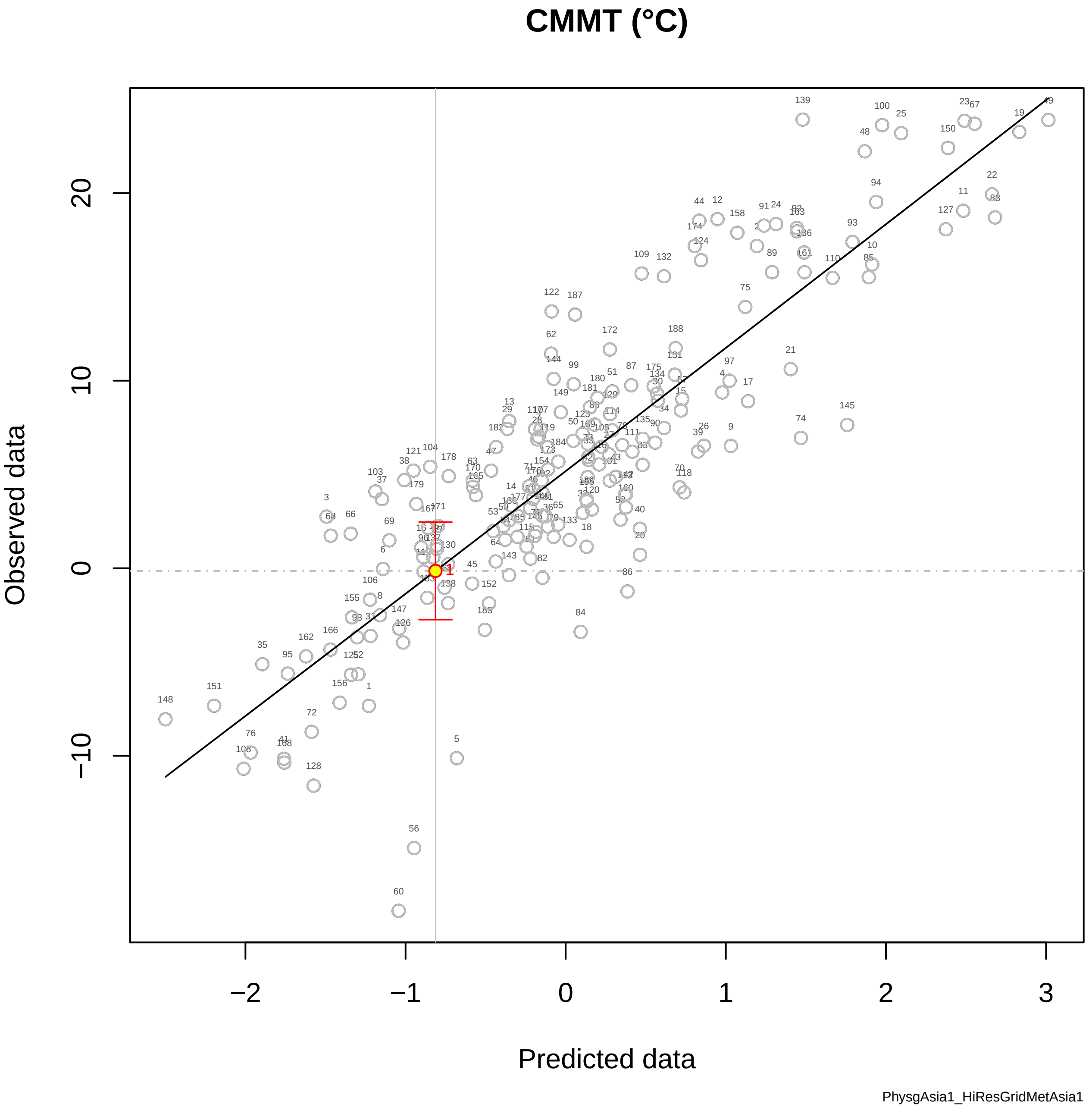

CMMT (°C)
Observed data
Predicted data
20
10
0
−10
−2
−1
0
1
2
3
PhysgAsia1_HiResGridMetAsia1

### GROWSEAS (months)

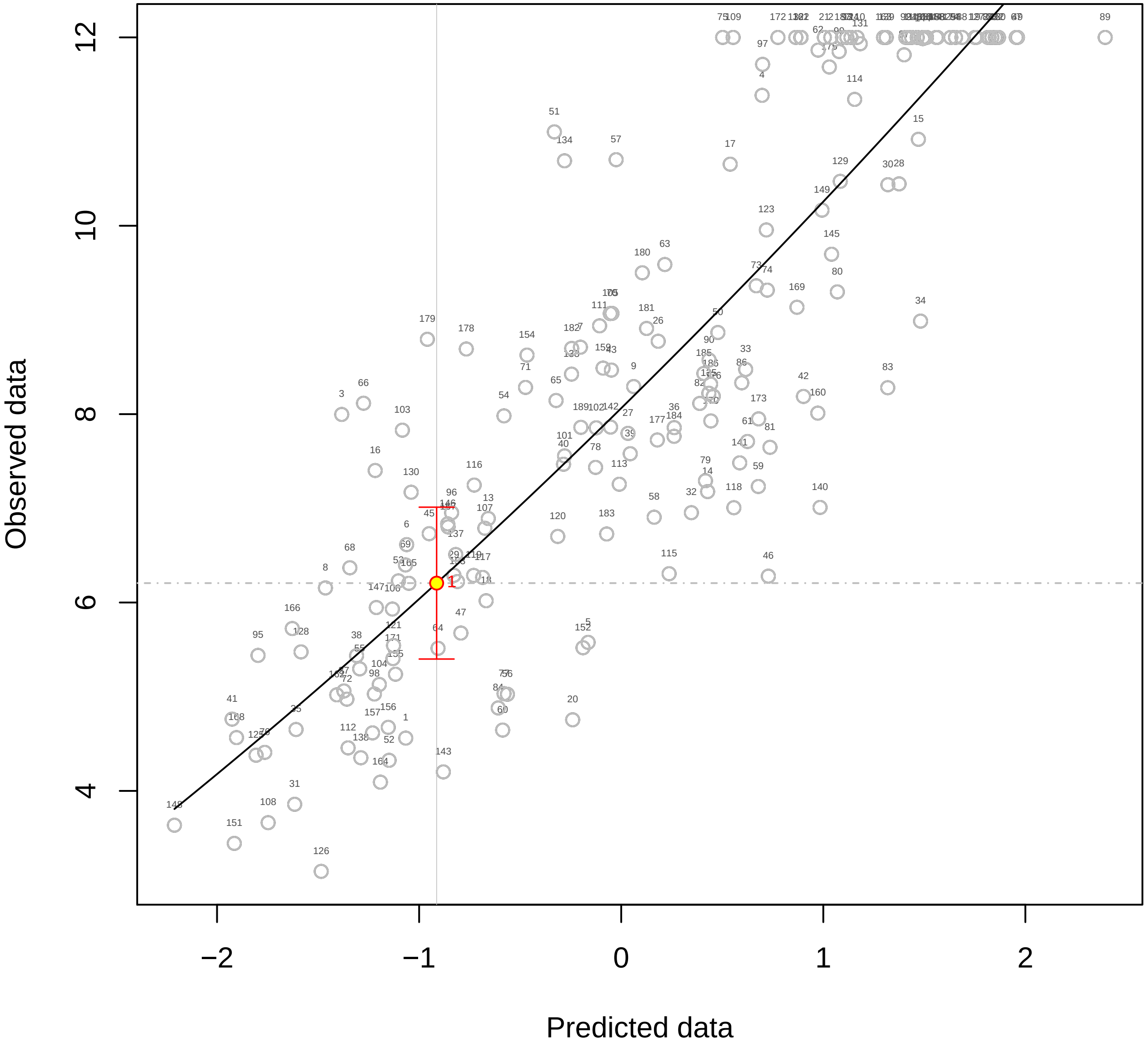


PhysgAsia1_HiResGridMetAsia1

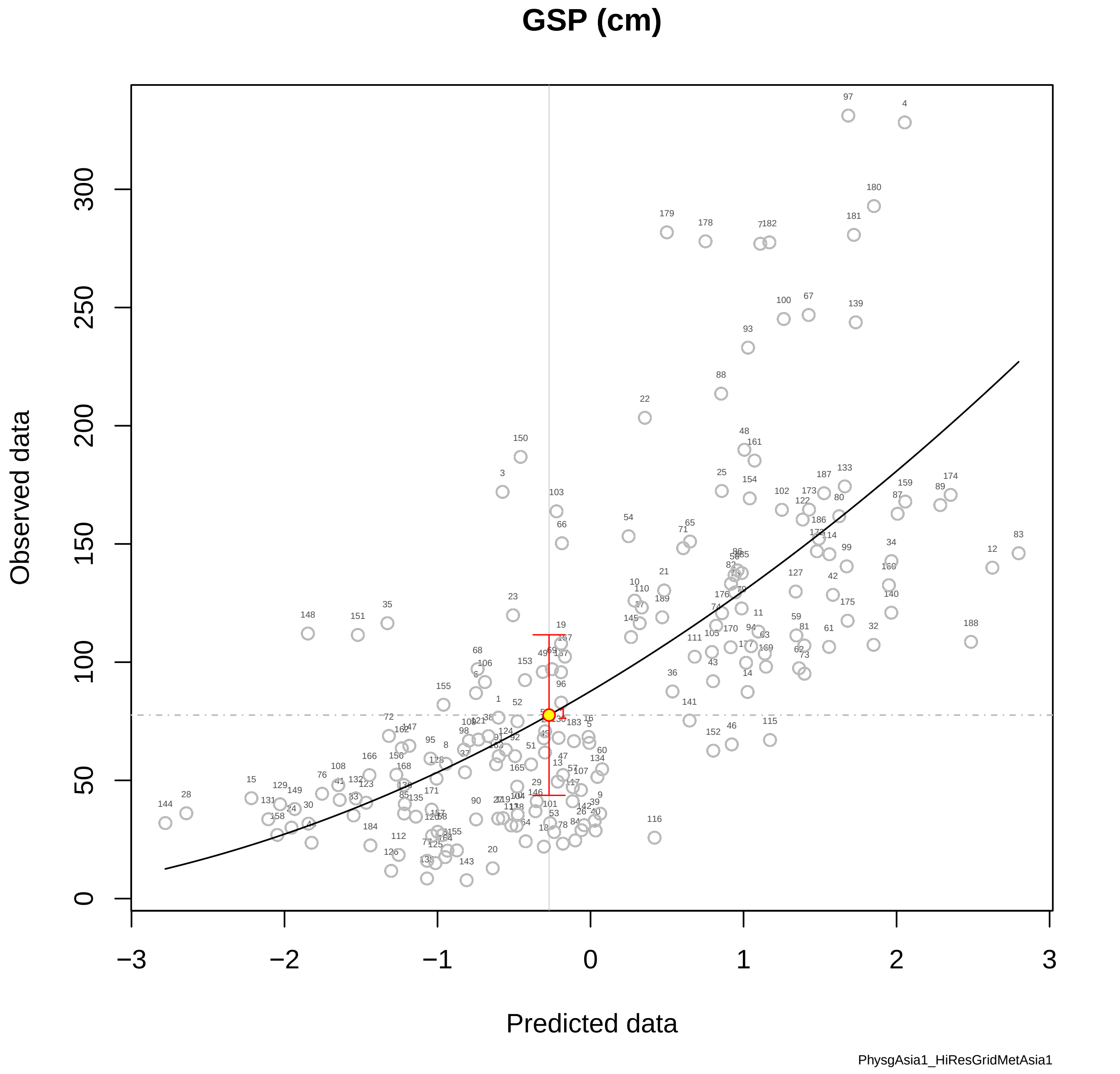

GSP (cm)
Observed data
Predicted data
0
50
100
150
200
250
300
−3
−2
−1
0
1
2
3
PhysgAsia1_HiResGridMetAsia1

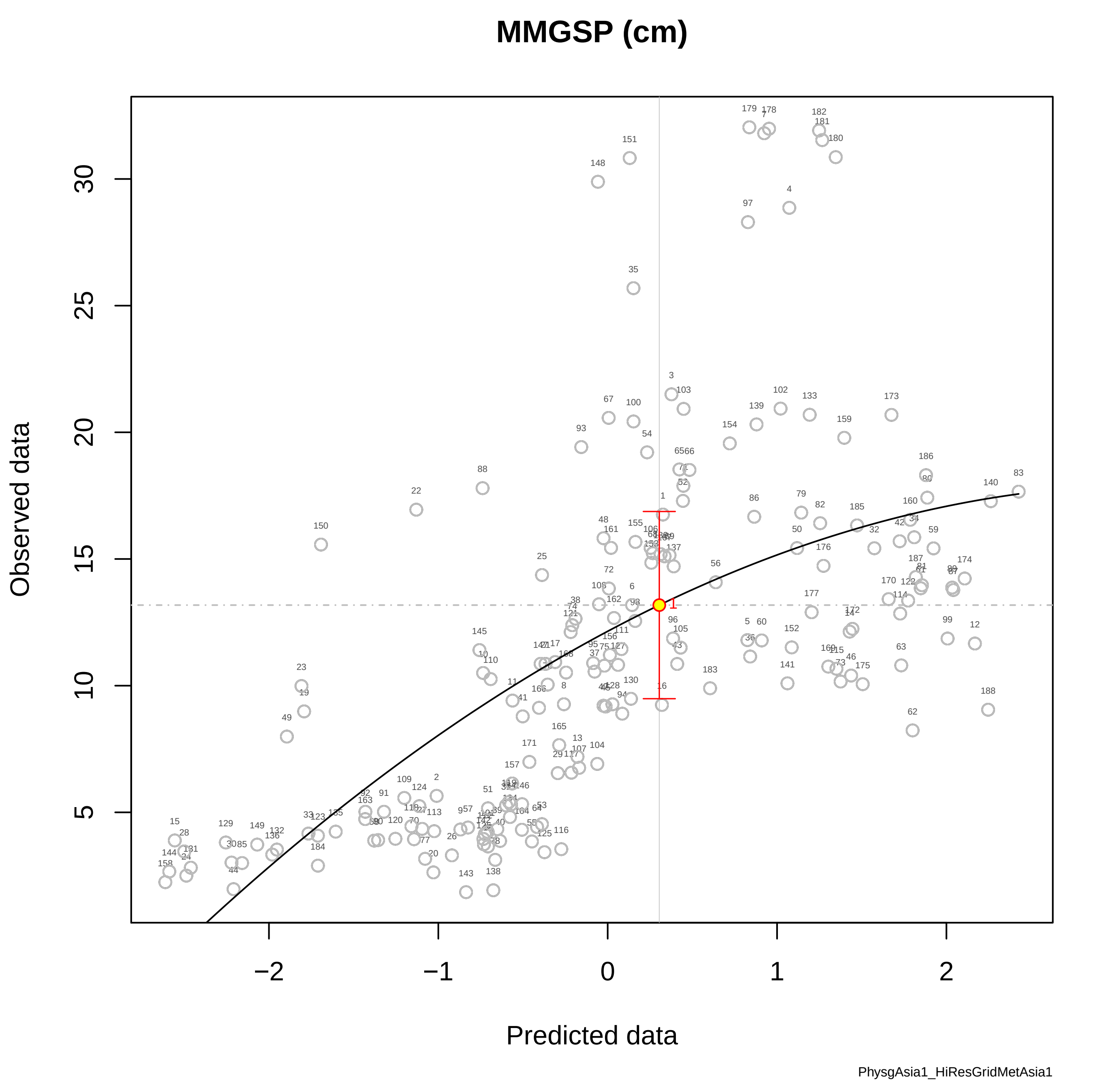
MMGSP (cm)
Observed data
Predicted data
PhysgAsia1_HiResGridMetAsia1

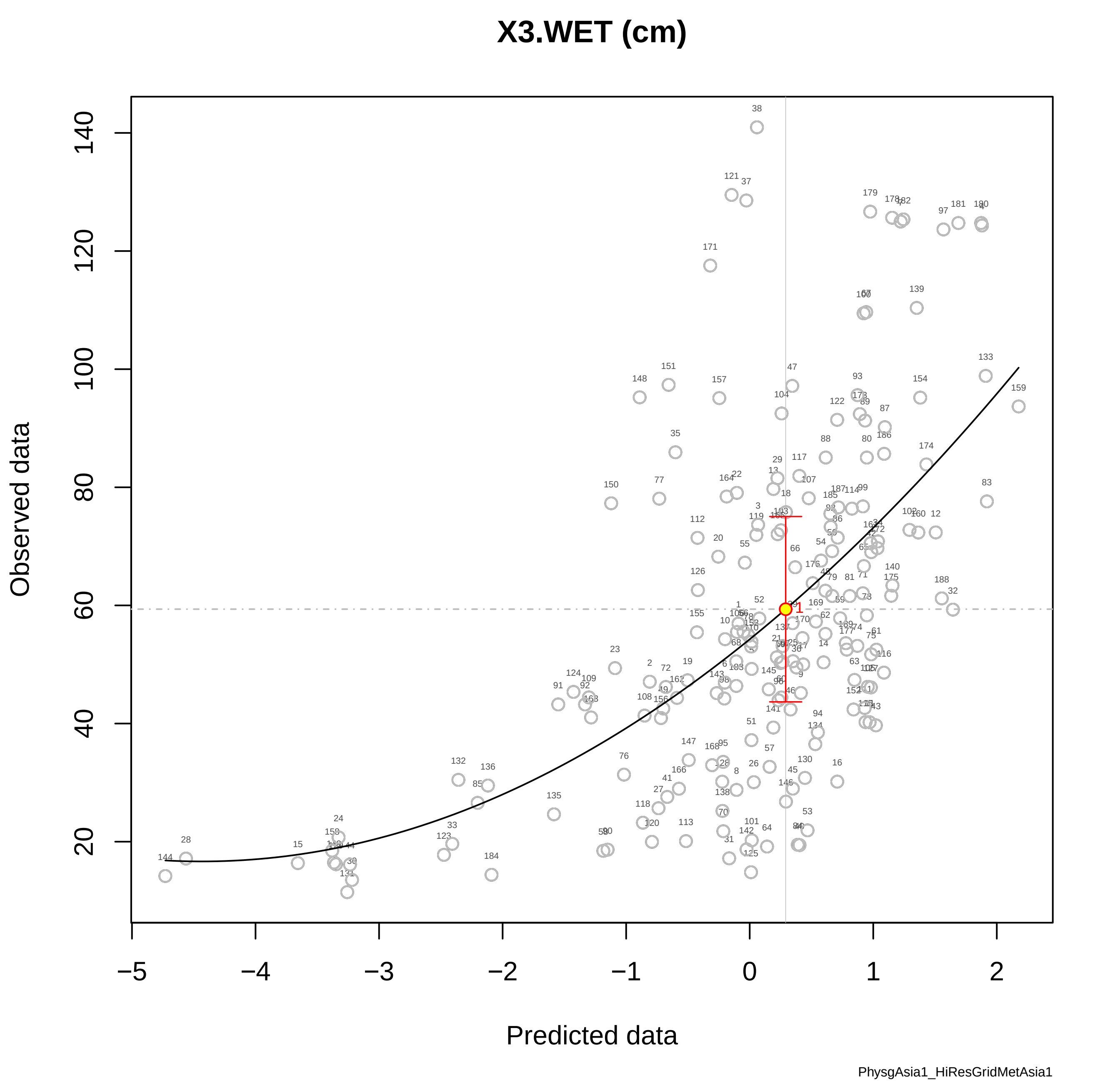
X3.WET (cm)
Observed data
Predicted data
PhysgAsia1_HiResGridMetAsia1

### X3.DRY (cm)

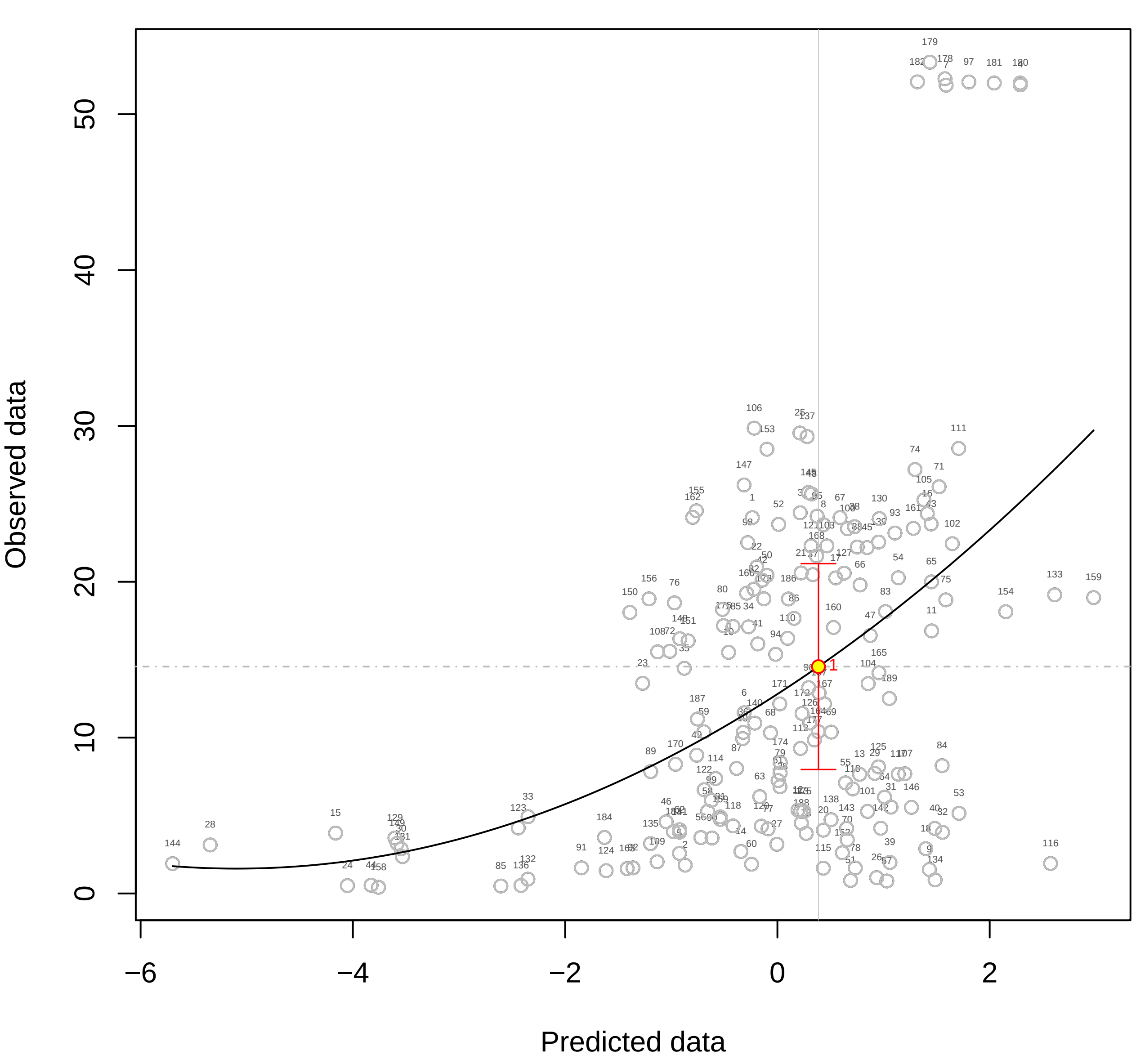


PhysgAsia1_HiResGridMetAsia1

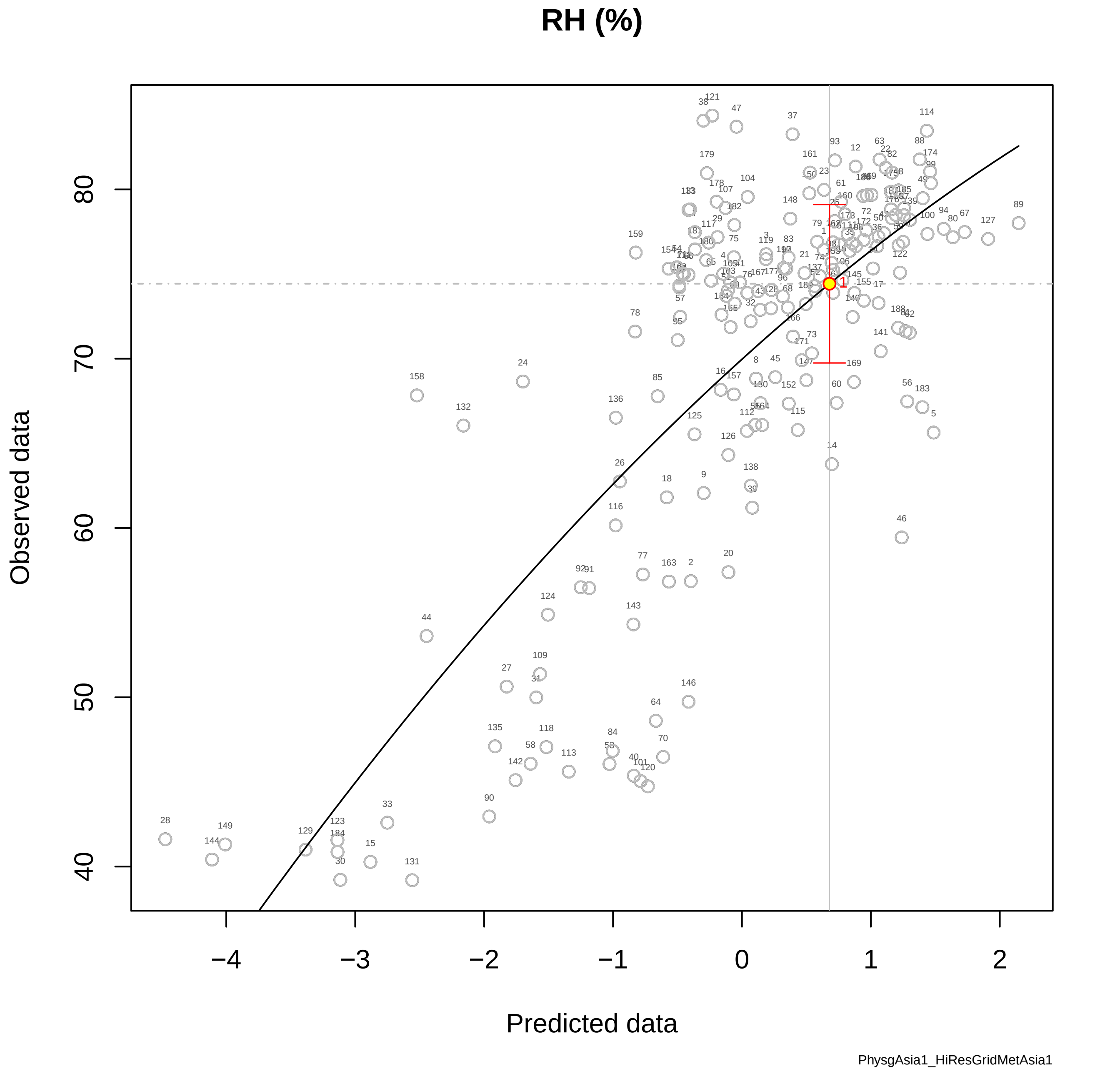

RH (%)
Observed data
Predicted data
PhysgAsia1_HiResGridMetAsia1

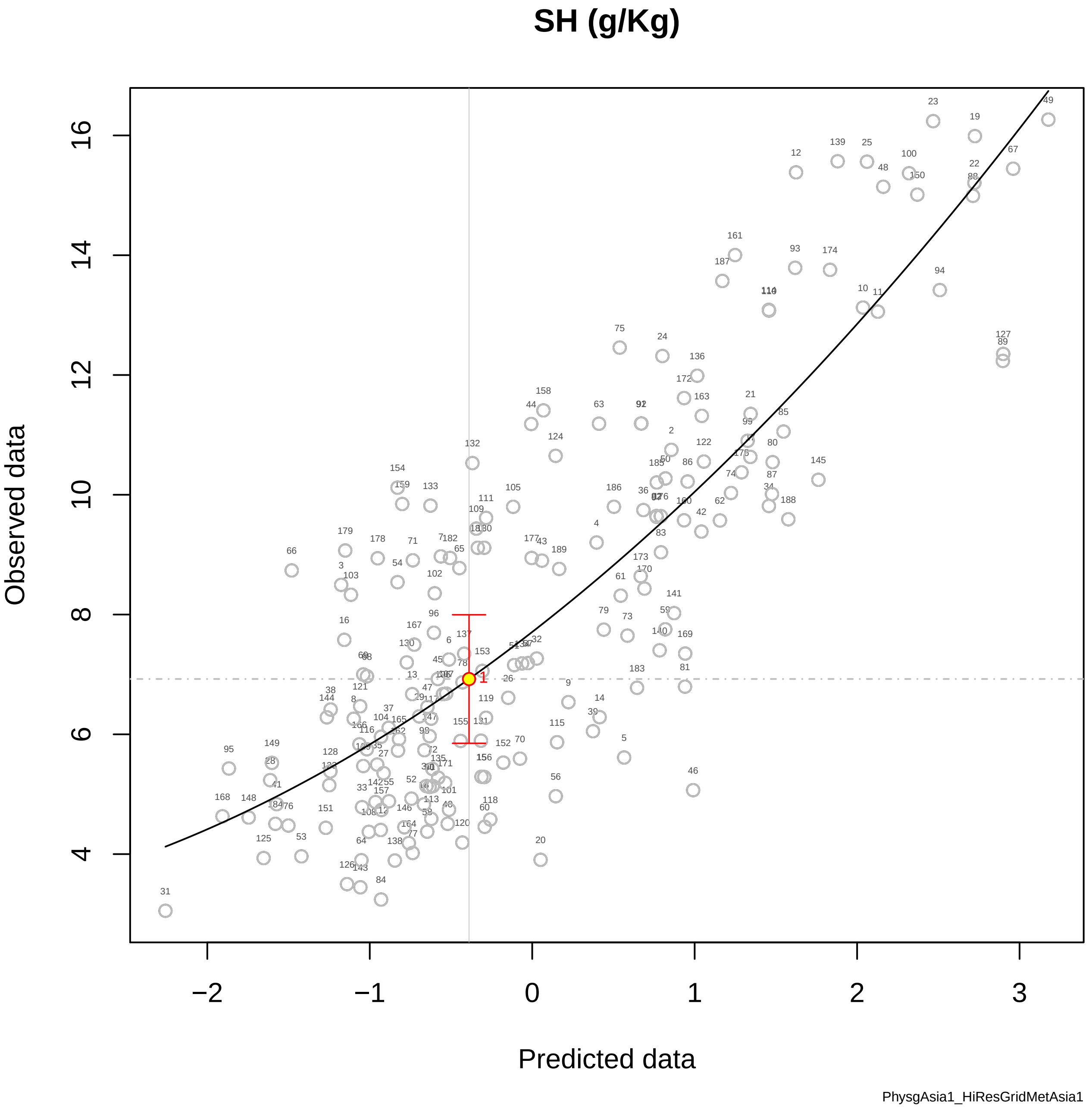

SH (g/Kg)
Observed data
Predicted data
PhysgAsia1_HiResGridMetAsia1

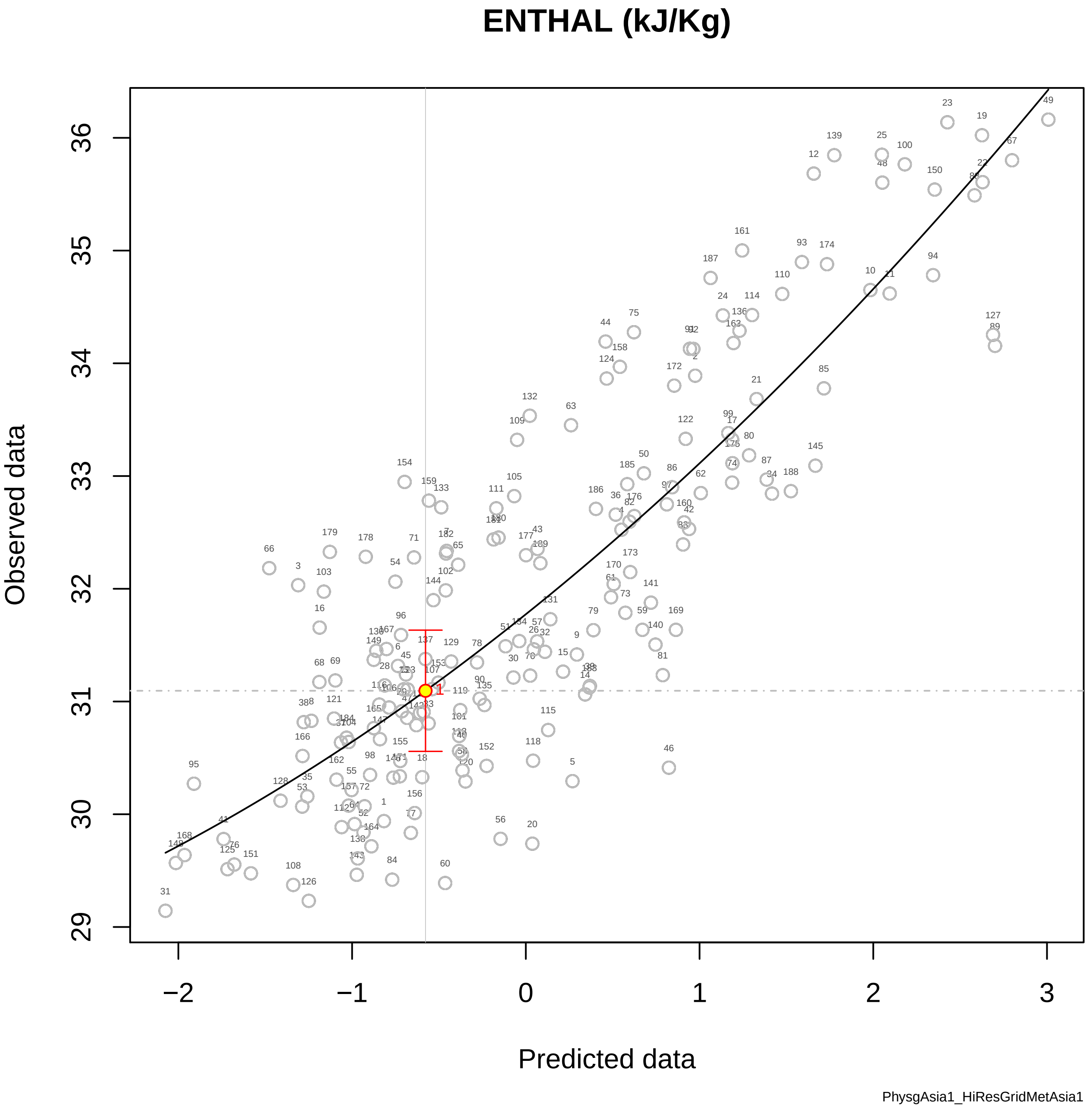
ENTHAL (kJ/Kg)
Observed data
Predicted data
29
30
31
32
33
34
35
36
−2
−1
0
1
2
3
PhysgAsia1_HiResGridMetAsia1
