## Supplementary figures and images for "Messinian vegetation and climate of the intermontane Florina-Ptolemais-Servia Basin, NW Greece inferred from palaeobotanical data: How well do plant fossils reflect past environments?" (from another version of this preprint: version 1, 2019-11-25)

### CCA1vsCCA2.jpg

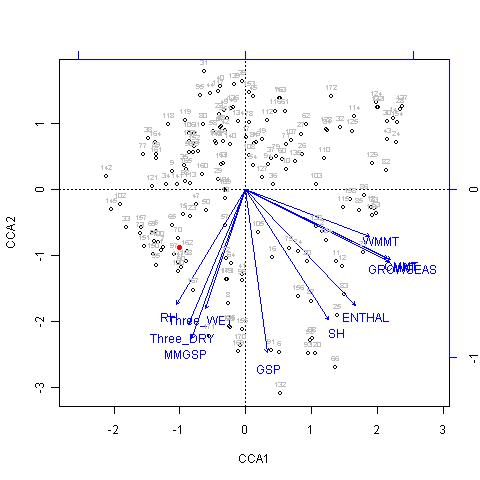

### CCA1vsCCA2.jpg

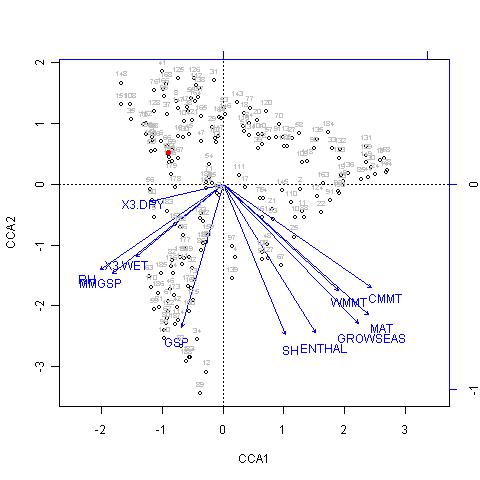

### CCA1vsCCA3.jpg

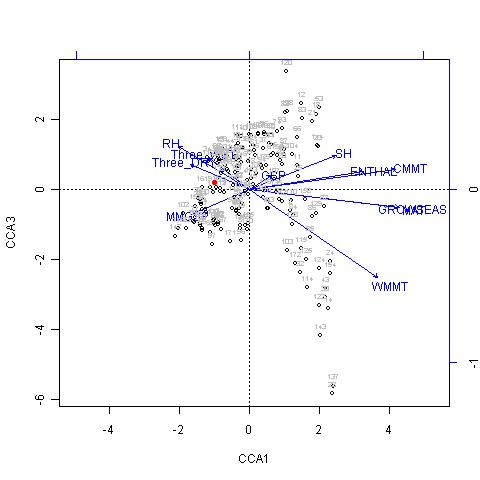

### CCA1vsCCA3.jpg

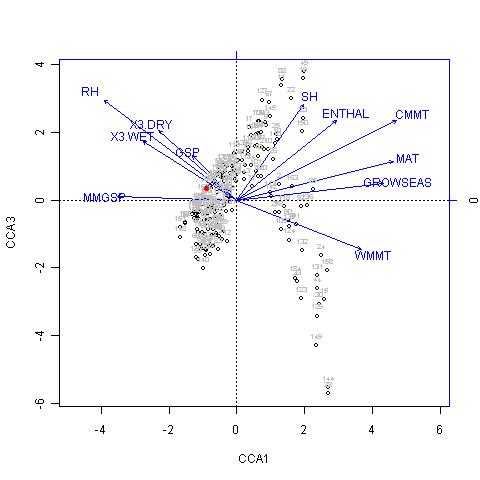

### CCA2vsCCA3.jpg

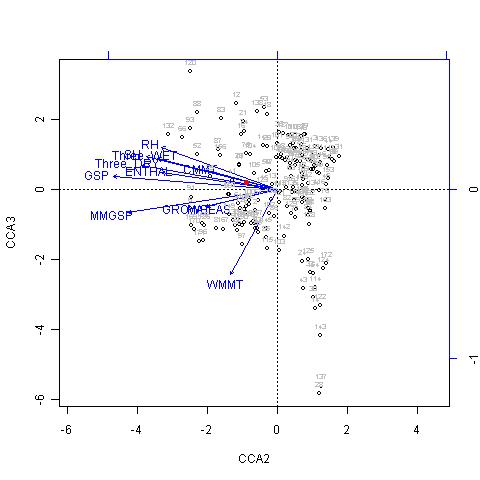

### CCA2vsCCA3.jpg

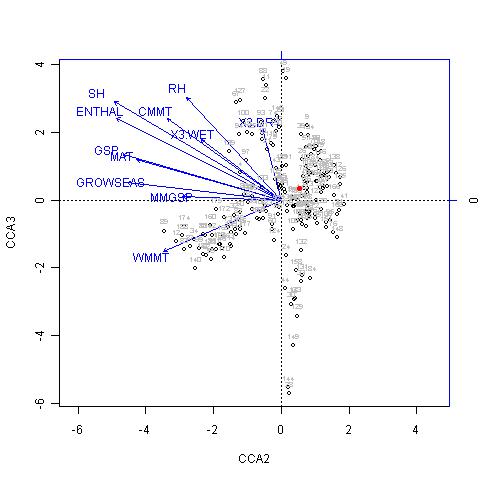

### CCA3d.jpg

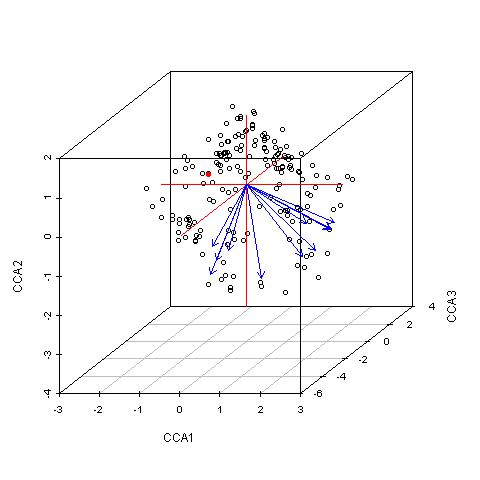

### CCA3d.jpg

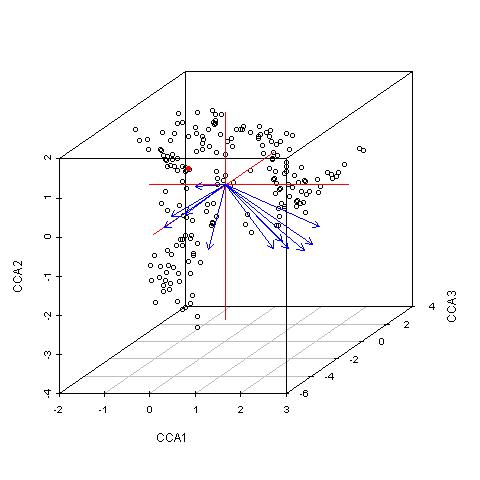

### CCAGraph.pdf

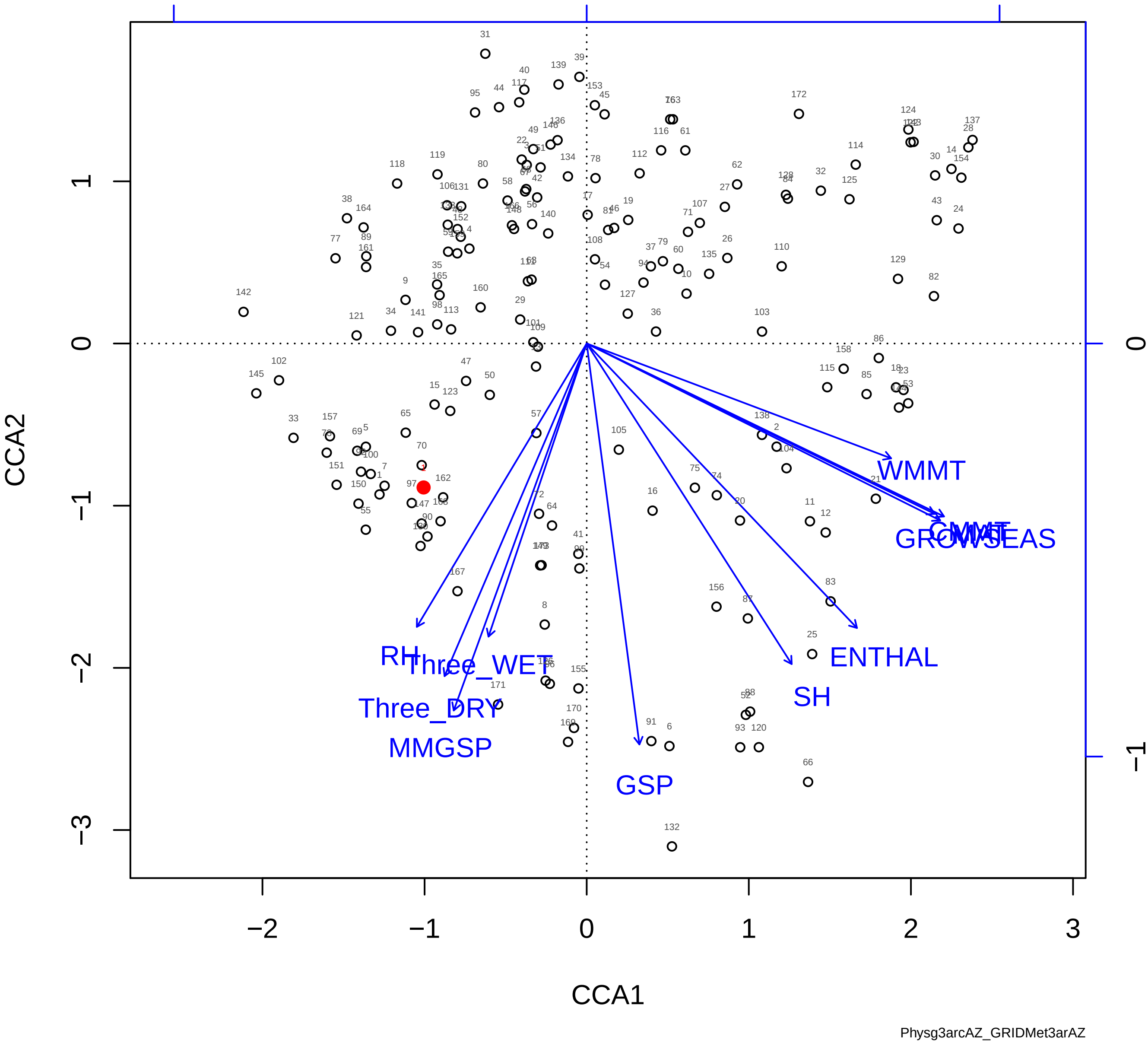

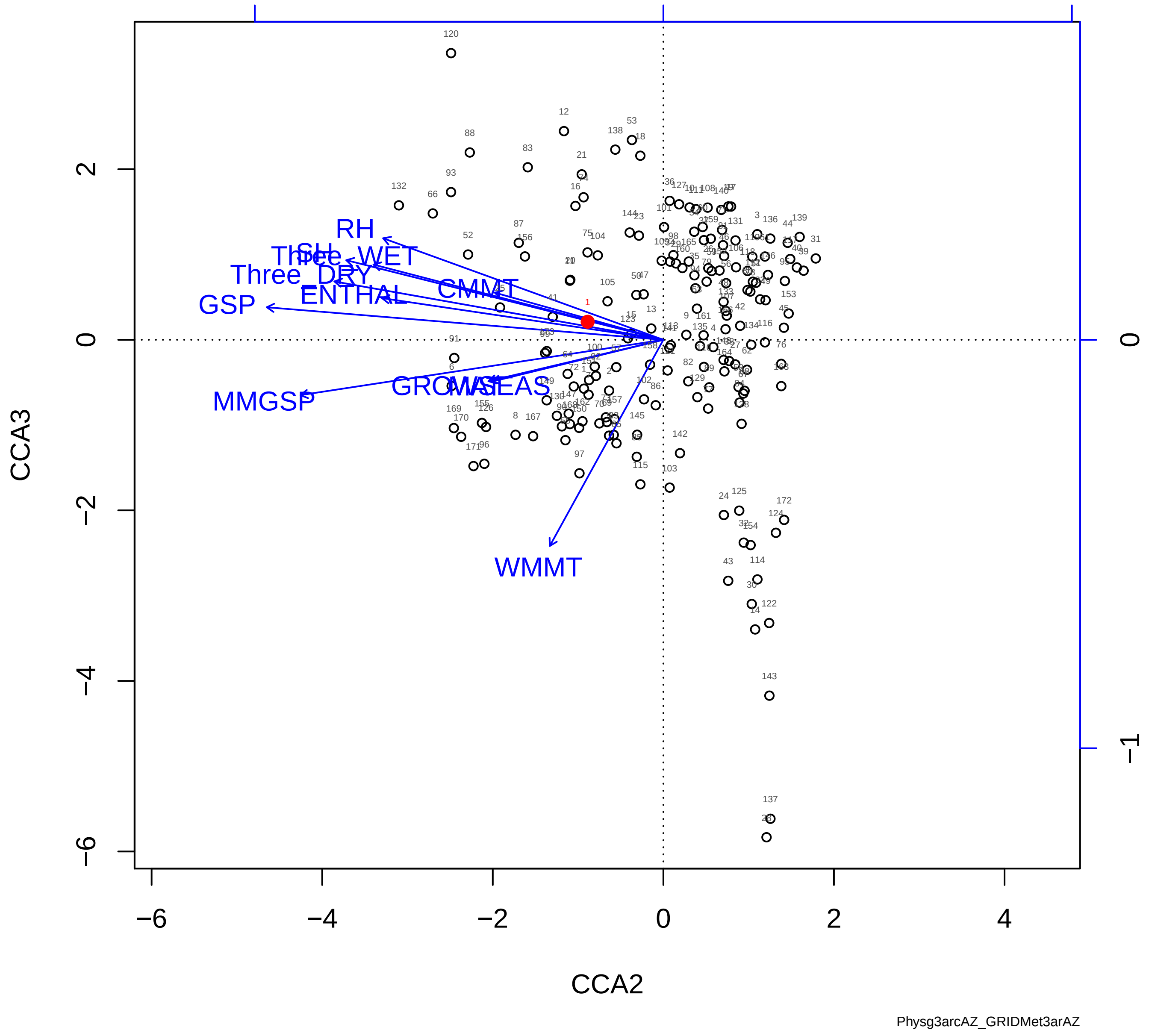

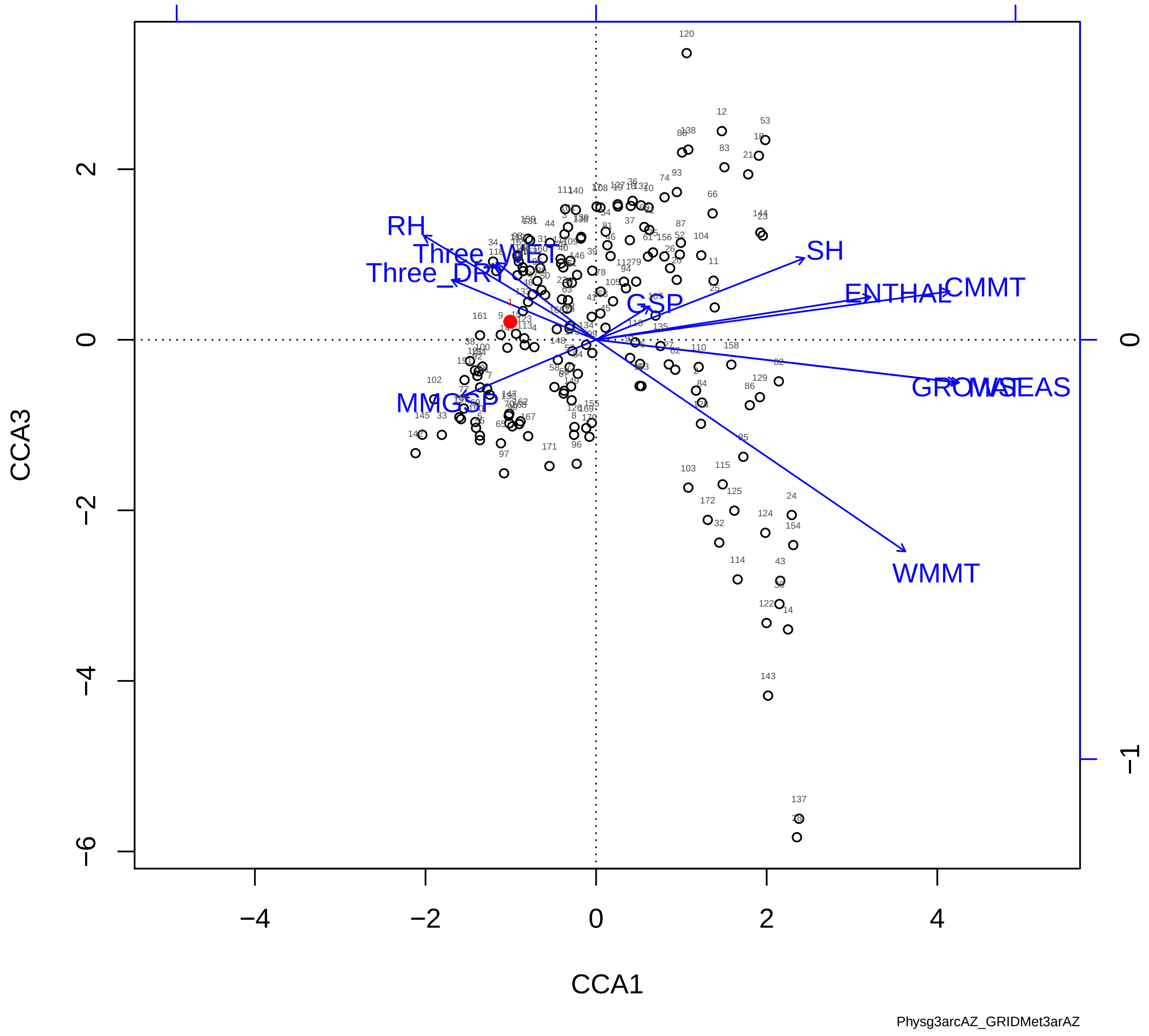

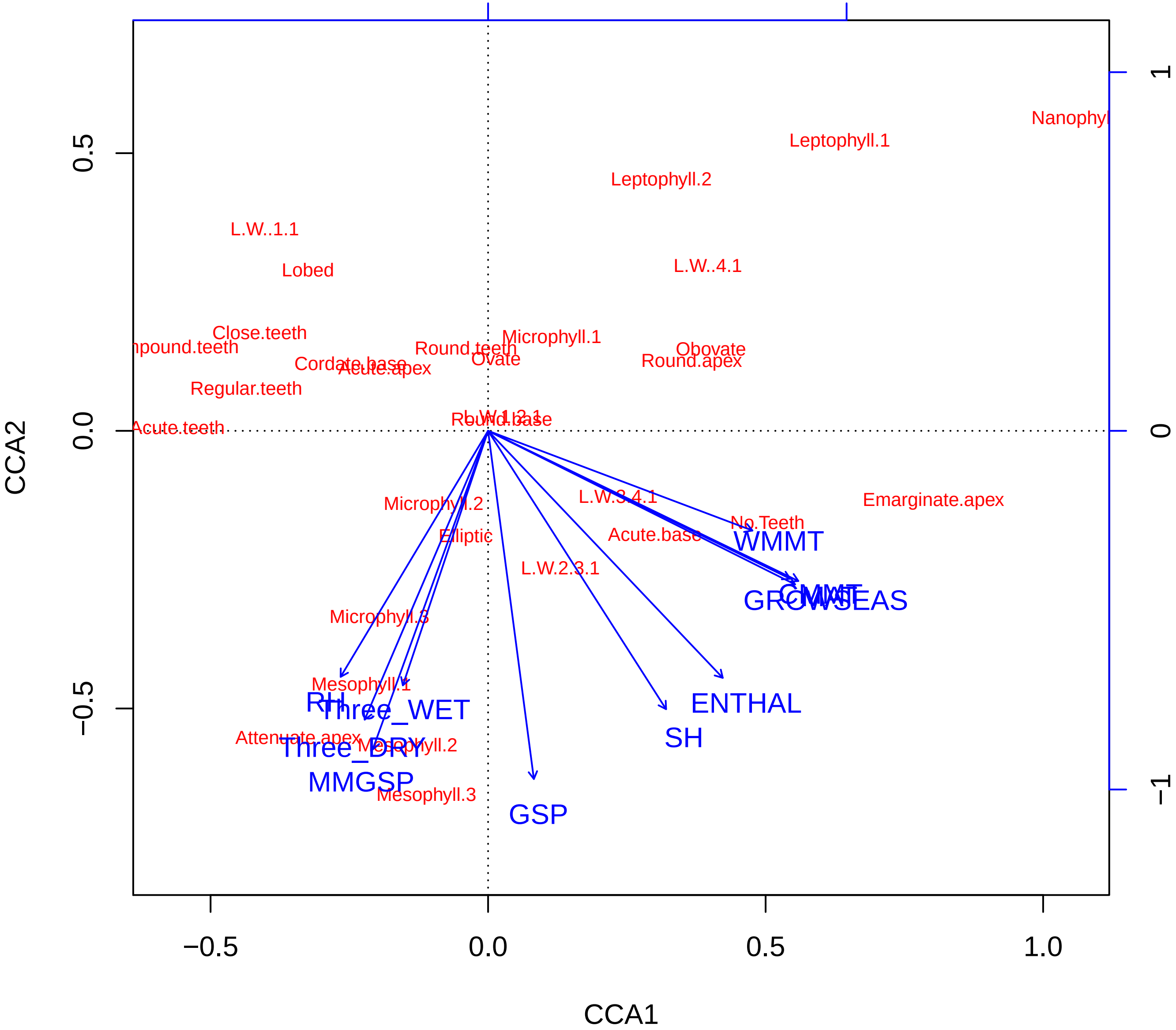

Physg3arcAZ_GRIDMet3arAZ

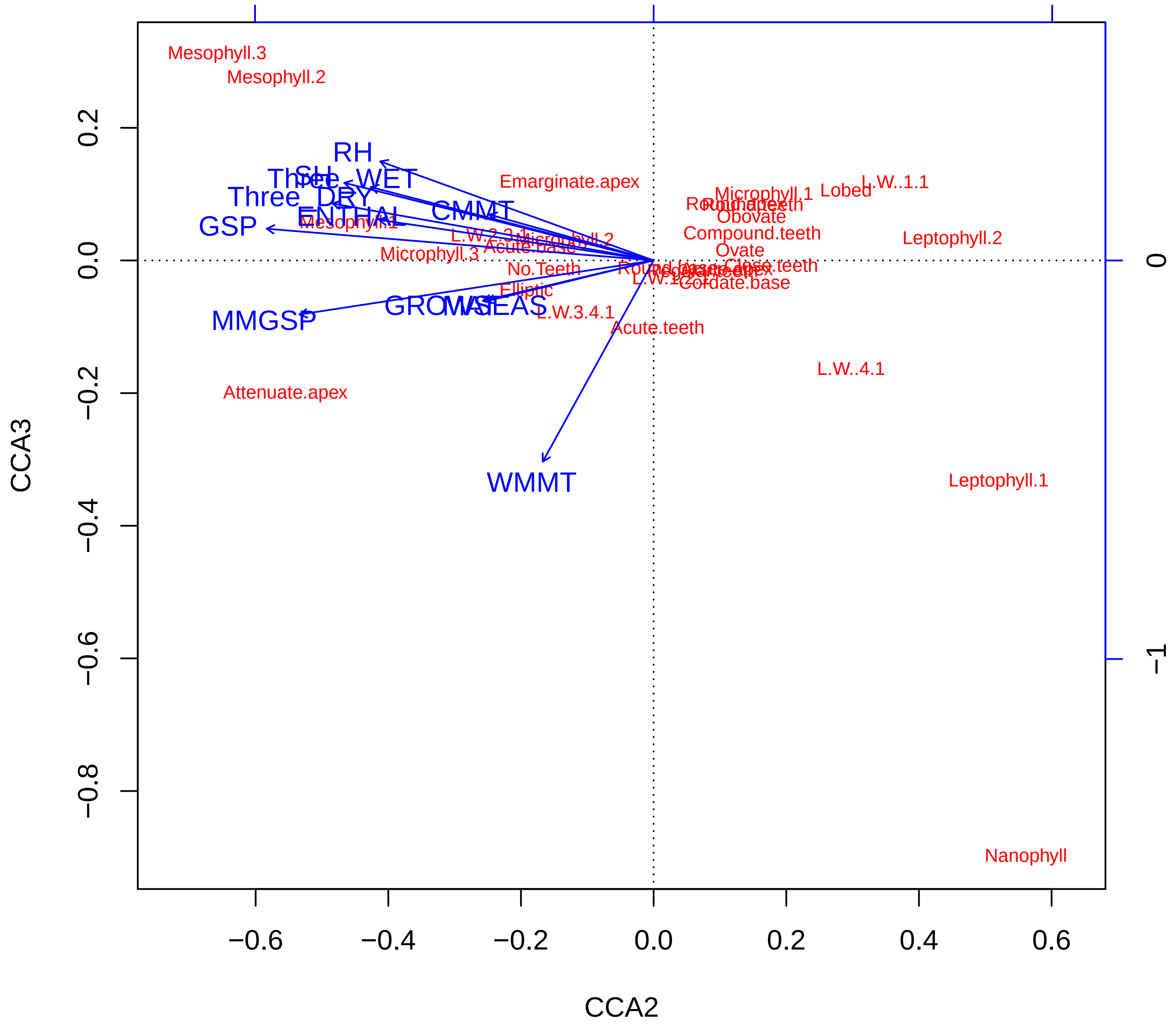


Physg3arcAZ_GRIDMet3arAZ

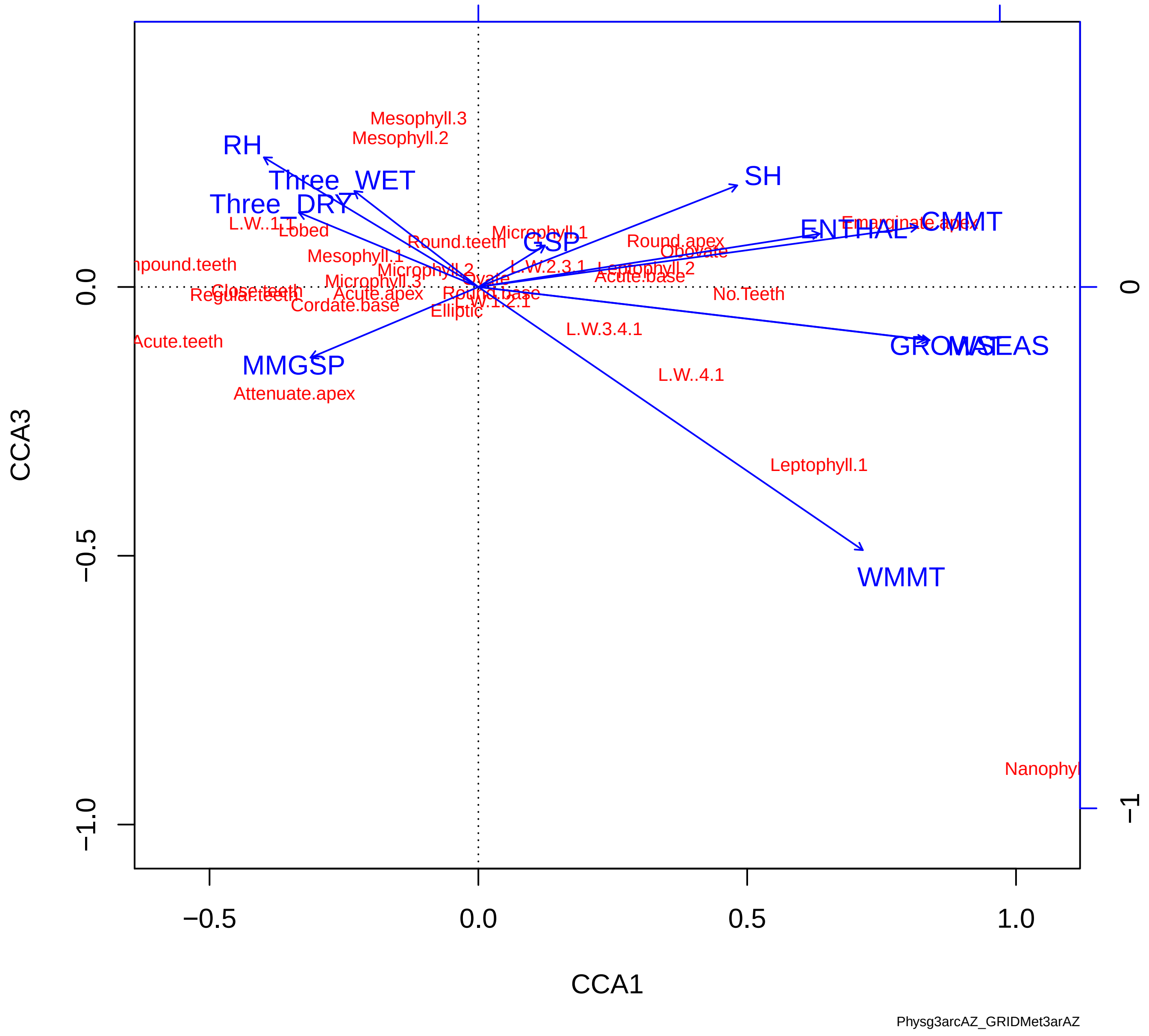

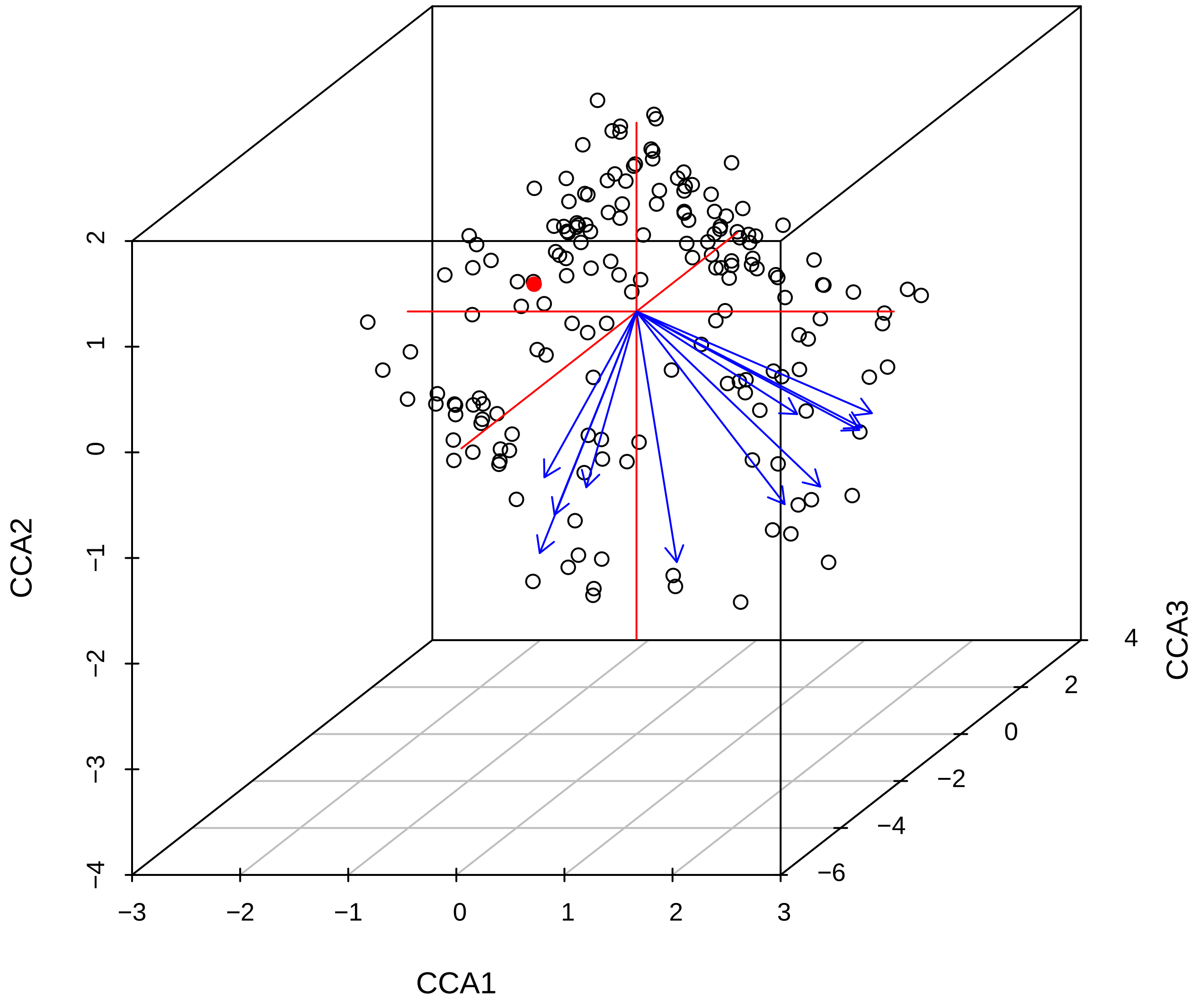


Physg3arcAZ_GRIDMet3arAZ

### GSP.jpg

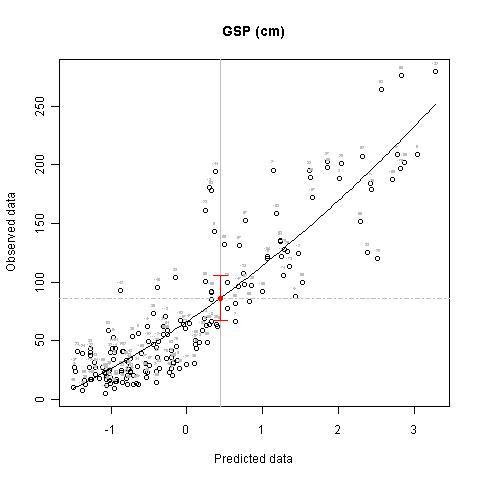

### GSP.jpg

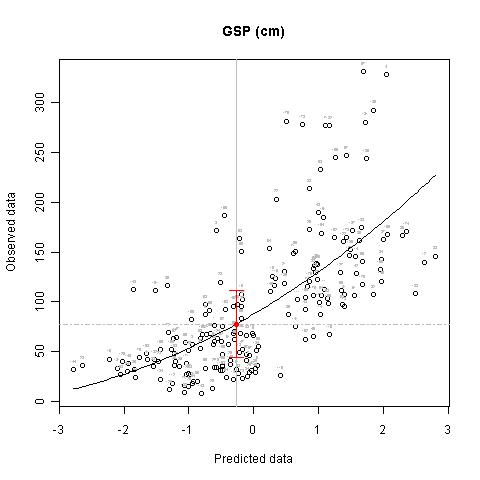

### MAT.jpg

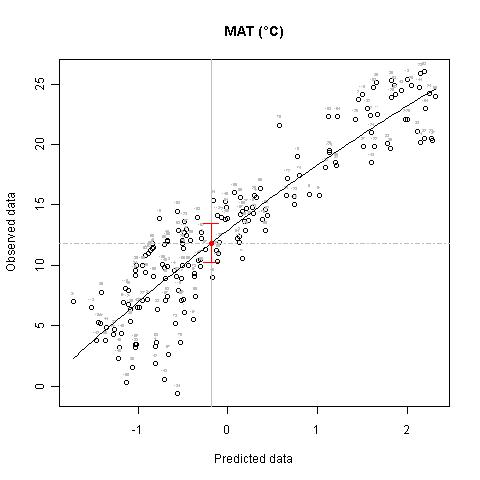

### MAT.jpg

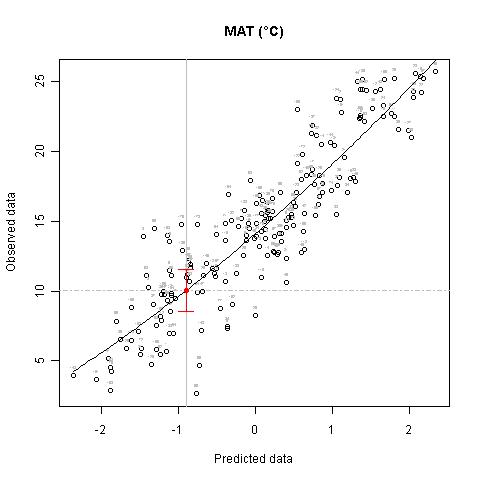
